# The visual white matter: The application of diffusion MRI and fiber tractography to vision science

**DOI:** 10.1101/072793

**Authors:** Ariel Rokem, Hiromasa Takemura, Andrew Bock, K. Suzanne Scherf, Marlene Behrmann, Brian Wandell, Ione Fine, Holly Bridge, Pestilli Franco

## Abstract

Visual neuroscience has traditionally focused much of its attention on understanding the response properties of neurons along the visual pathways. This review focuses instead on the properties of the white matter connections between these neurons. Specifically, we provide an introduction to methods to study the human visual white matter using diffusion MRI (dMRI). This method allows us to measure the white matter connections in individual visual systems in vivo, allows us to trace long-range connections between different parts of the visual system, and to measure the biophysical properties of these connections. We explain the principles underlying dMRI measurements and the basics of modeling these data. We review a range of findings from recent studies on connections between different visual field maps, on the effects of visual impairment on the white matter, and on the properties underlying networks that process visual information that supports visual face recognition. Finally, we discuss a few promising directions for future studies. These include new methods for analysis of MRI data, open data-sets that are becoming available to study brain connectivity and white matter properties, and open-source software for the analysis of these data.

## 1 Introduction

The cerebral hemispheres of the human brain can be subdivided into two primary tissue types: the white matter and the gray matter (Fields, 2008a). Whereas the gray matter contains neuronal cell bodies, the white matter contains primarily the axons of these neurons, and glial cells. The axons constitute the connections that transmit information between distal parts of the brain - *mm* to *cm* in length. Some types of glia (primarily oligodendrocytes) form an insulating layer around these axons (myelin sheaths) that allow the axons to transmit information more rapidly (Waxman and Bennett, 1972), and more accurately (Kim et al., 2013), and reduce their energy consumption (Hartline and Colman, 2007).

Much of neuroscience has historically focused on understanding the functional response properties of individual neurons and cortical regions (Fields, 2013, 2004). It has often been implicitly assumed that white matter, and the long-range neuronal connections it supports have a binary nature: either they are connected and functioning, or they are disconnected. However, more recently there is an increasing focus on the role that the variety of properties of the white matter may play in neural computation (Bullock et al., 2005; Jbabdi et al., 2015; Reid, 2012; Sporns et al., 2005) together with a growing understanding of the importance of the brain networks composed of these connections in cognitive function (Petersen and Sporns, 2015).

The renewed interest in white matter is in part due to the introduction of new technologies. Long-range connectivity between parts of the brain can be measured in a variety of ways. For example, functional magnetic resonance imaging (fMRI) is used to measure correlations between the blood-oxygenation level dependent (BOLD) signal in different parts of the brain. In this review, we focus on results from diffusion-weighted Magnetic Resonance Imaging technology (dMRI). Together with computational tractography dMRI provides the first opportunity to measure white matter and the properties of long-range connections in the living human brain. We will refer to these connections as “fascicles”, an anatomical term that refers to bundles of nerve or muscle. Measurements in living brains demonstrate the importance of white-matter for human behavior, health and disease (Fields, 2008a; Jbabdi et al., 2015; Johansen-Berg and Behrens, 2009) development (Lebel et al., 2012; Yeatman et al., 2014a) and learning (Bengtsson et al., 2005; Blumenfeld-Katzir et al., 2011; Hofstetter et al., 2013; Sagi et al., 2012; Sampaio-Baptista et al., 2013; Thomas and Baker, 2013). The white matter comprises a set of active fascicles that respond to human behavior, in health and disease by adapting their density, their shape, and their molecular composition and correspond to human cognitive, motor and perceptual abilities.

The visual white matter sustains visual function. In the primary visual pathways, action potentials travel from the retina via the optic nerve, partially cross at the optic chiasm, and continue via the optic tract to the lateral geniculate nucleus (LGN) of the thalamus. From the LGN, axons carry visual information laterally through the temporal lobe, forming the structure known as Meyer’s loop, and continue to the primary visual cortex (Figure 1). The length of axons from the retina to the LGN is approximately 5 cm long, while the length of the optic radiation from the anterior tip of Meyer’s loop to the calcarine sulcus is approximately 10 cm, with an approximate width of 2 cm at the lateral horn of the ventricles (Peltier et al., 2006; Sherbondy et al., 2008b). Despite traveling the entire length of the brain, responses to visual stimuli in cortex arise very rapidly (Maunsell and Gibson, 1992), owing to the high degree of myelination of the axons in the primary visual pathways. Healthy white matter is of crucial importance in these pathways, as fast and reliable communication of visual input is fundamental for healthy visual function

**Figure 1:**
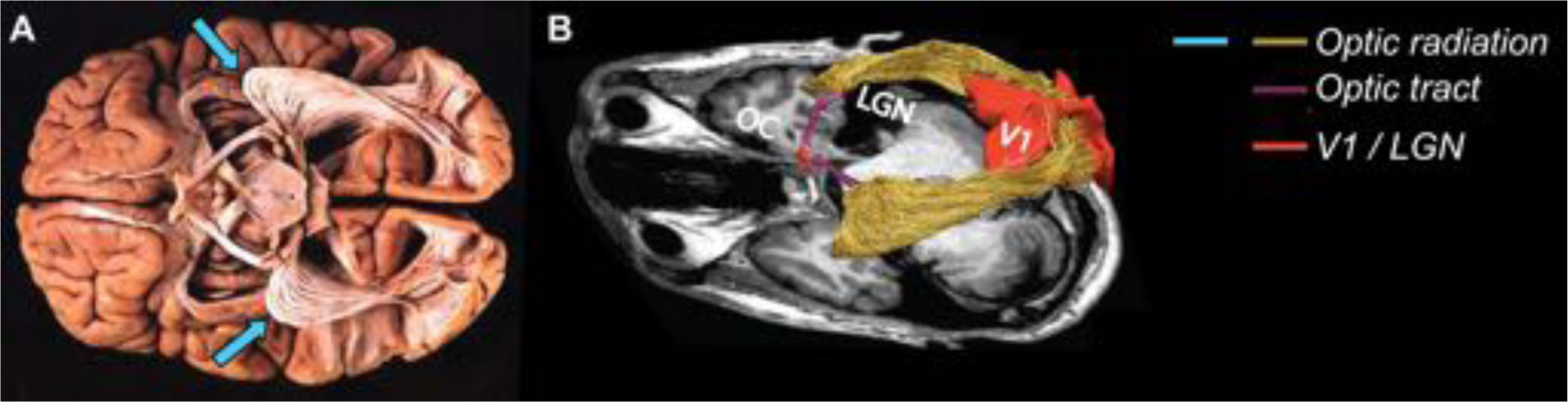
Postmortem and in vivo study of the visual white matter. **A**. Post mortem dissection of the optic radiation (OR), showing the Meyer’s loop bilaterally (blue arrows; adapted from (Sherbondy et al., 2008b). **B**. In vivo dissection of the optic radiation showing the optic chiasm (OC), Meyer’s loop, the optic tract as well as both primary visual cortex (V1) and the lateral geniculate nucleus (LGN; (Ogawa et al., 2014))

**Figure 2:**
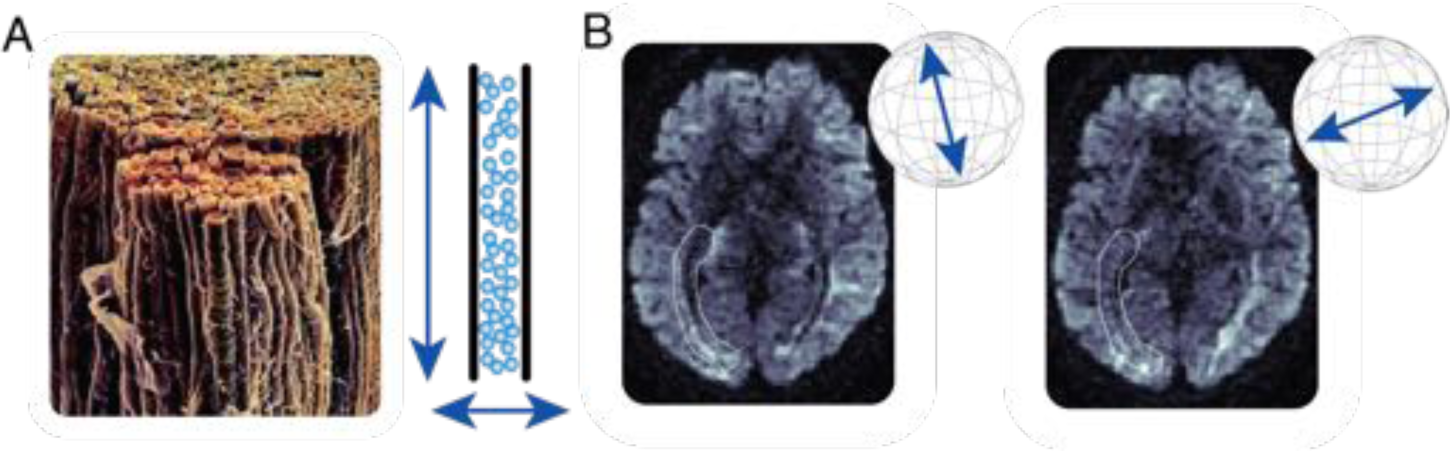
Inferences about white matter from measurements of water diffusion. **A** Micro-graph of the human optic nerve. Left, bundles of myelinated axonal fascicles from the human optic nerve (image courtesy of Dr. George Bartzokis). Right, cartoon example of anisotropic water diffusion restriction by white matter fascicles. Water diffuses further along the direction of the “tube” (Pierpaoli and Basser, 1996). **B** Example of diffusion-weighted magnetic resonance measurements in a single slice of a living human brain. The same brain slice is shown as imaged with two different diffusion weighting directions. Diffusion weighting directions are shown by the inset arrows. The white highlighted area indicates the approximate location of part of the OR. The longitudinal shape of the OR appears as a local darkening of the MRI image when the diffusion weighting gradient is aligned with the direction of the myelinated fascicles within the OR, right-hand panel

This review presents the state of the art in dMRI studies of the human visual system. We start with an introduction of the measurements and the analysis methods. Following that, we review three major applications of dMRI:

1. The delineation of the relationship between visual field maps and the white matter connections between the maps. This section focuses on a test-case of the connections between dorsal and ventral visual field maps.
2. The effects of disorders of the visual system on the visual white matter. This section focuses on the changes that occur in the white matter in cases of peripheral and cortical blindness, and on white matter connectivity that underlies cross-modal plasticity and residual vision respectively in these cases.
3. The role that the visual white matter plays in visual discrimination of specific categories of visual objects. In particular, this section focuses on the white matter substrates of the perception of face stimuli.

Taken together, these three sections present multiple facets of dMRI research in vision science, covering a wide array of approaches and applications. They all demonstrate the ways in which studying the white matter leads to a more complete understanding of the biology of the visual system: its structure, response properties and relation to perception and behavior. In the summary of the review we will point out some common threads among these applications, point to a few of the challenges facing the field, and the promise of future developments to help address some of these challenges.

## Methods for in-vivo study of the human white matter

In this section, we introduce concepts and methods for measuring white matter connections using dMRI. The text is accompanied by a series of code examples implemented as Jupyter notebooks, which can be downloaded from https://github.com/arokem/visual-white-matter.

### Measuring white matter using diffusion magnetic resonance imaging

Diffusion MRI measures the directional diffusion of water in different locations in the brain. In MRI, a strong magnetic field is applied to the brain, causing protons in water molecules to precess at a field-strength-dependent frequency. When the magnitude of the magnetic field is varied linearly in space (creating a a magnetic field gradient), protons in different locations along the gradient begin to precess at different rates. The rate of precession varies spatially along the same direction as the magnetic gradient. Diffusion MRI utilizes a second gradient pulse with the same magnitude but in the opposite direction that refocuses the signal by aligning the spins of water molecules. Refocusing will not be perfect for protons that have moved during the time interval between the gradient pulses, and the resulting loss of signal from dephasing is used to infer the mean diffusion of water molecules (due to random Brownian motion) along the direction of the magnetic gradient. Measurements conducted with gradients applied along different axes sensitize the signal to diffusion in different directions (Figure 1B).

Additional magnetic field gradients are used to render the measurements spatially selective. In-vivo measurements in human typically use resolutions on the order of 2 × 2 × 2 *mm*^3^ (isotropic voxels). Using modern imaging techniques, such as multi-band imaging (Feinberg et al., 2010), high SNR measurements can be done in vivo with voxel sizes as small as 1.25-1.5 *mm* (isotropic) (Setsompop et al., 2013) using standard 3 Tesla (3T) hardware. Post-mortem and animal measurements can be done with much smaller size scale, down to approximately 200 *µm* (isotropic) (Leuze et al., 2014; Reveley et al., 2015).

The sensitivity of the measurement to diffusion increases with the amplitude and duration of the magnetic field gradient, as well as with the time between the two gradient pulses (the *diffusion time* or *mixing time*). These three parameters are usually summarized in a single number: the b-value (see code example: http://arokem.github.io/visual-white-matter/signal-formation). Increased diffusion-sensitivity does come at a cost: measurements at higher b-values also have lower SNR. In the typical dMRI experiment, diffusion times are on the order of 50-100 *msec*. Within this time-scale, at body temperature, water molecules may diffuse an average of approximately 20 *µm*, when no restriction is placed on their diffusion (at body temperature) (Le Bihan and Iima, 2015). This rate of diffusion is found in the ventricles of the brain, for example. Within other brain structures the average diffusion distance is impeded by the brain tissue (Figure 1; see also: http://arokem.github.io/visual-white-matter/dMRI-signals).

In regions of the brain or body that contain primarily spherically-shaped cell bodies, water diffusion is equally restricted in all directions: it is isotropic. On the other hand, in fibrous biological structures, such as muscle or nerve fascicles, the diffusion of water is restricted across the fascicles (e.g. across axonal membranes) more than along the length of the fascicles (e.g. within the axoplasm). In these locations, diffusion is anisotropic. Measurements of isotropic and anisotropic diffusion can be used to estimate the microstructural properties of brain tissue and brain connectivity.

Inferring brain tissue properties and connectivity from dMRI data generally involves two stages: the first estimates the microstructure of the tissue locally in every voxel. The second stage connects local estimates across voxels to create a macrostructural model of the axonal pathways. We review approaches to these two stages in the sections that follow.

### Estimating brain microstructure: models of diffusion in single voxels

Even though diffusion MRI samples the brain using voxels at millimeter resolution, the measurement is sensitive to diffusion of water within these voxels at the scale of *µm*. This makes dMRI a potent probe of the aggregate microstructure of the brain tissue in each voxel. A variety of analytic techniques and different models are routinely used to model tissue microstructure in brain voxel. We present two of the major approaches below.

#### The diffusion tensor model

The diffusion tensor model (DTM; (Basser et al., 1994a,b); see: http://arokem.github.io/visual-white-matter/dtm) approximates the directional profile of water diffusion in each voxel as a 3D Gaussian distribution. The direction in which this distribution has its largest variance is an estimate of the principal diffusion direction (PDD; Figure 3) – the direction of maximal diffusion. In many places in the brain, the PDD is aligned with the orientation of the main population of nerve fascicles and can be used as a cue for tractography (see below). In addition to the PDD, the diffusion tensor model provides other statistics of diffusion: the mean diffusivity (MD), which is an estimate of the average diffusivity in all directions, and the fractional anisotropy (FA), which is an estimate of the variance in diffusion in different direction (Pierpaoli and Basser, 1996) (Figure 3). These statistics are useful for characterizing the properties of white matter and numerous studies have shown that variance in these statistics within relevant white matter tracts can predict individual differences in perception (Genç et al., 2011), behavioral and cognitive abilities (e.g. (Yeatman et al., 2011), and mental health (Thomason and Thompson, 2011). A major advantage of the DTM is that it can be estimated from relatively few measurement directions: accurate (Rokem et al., 2015) and reliable (Jones, 2004) estimates of the tensor parameters require measurements in approximately 30-40 directions, which requires approximately 10 minutes for a full-brain scan in a clinical scanner.

**Figure 3:**
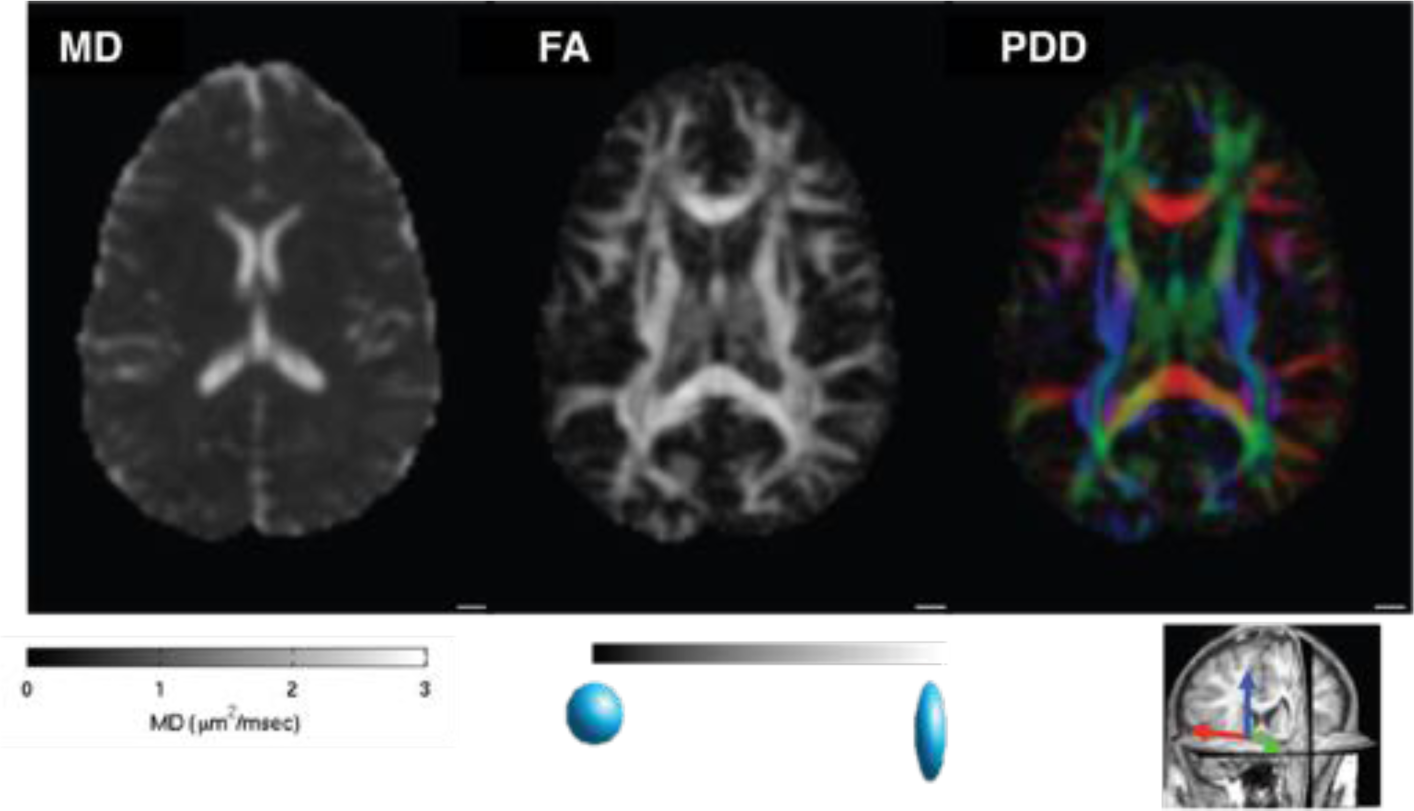
Diffusion tensor imaging. The figure shows the three primary estimates of white matter organization derived from the diffusion tensor model measured in a single slice of a human brain (spatial resolution = 2 *mm*^3^, b-value = 900*s/mm*^2^). **Mean diffusivity** (MD, left) represents water diffusion averaged across all measured diffusion direction. The value is expected to be 3 *µm*^2^/*msec* for pure water (in body temperature) and indeeed, MD is close to 3 in the ventricles. **Fractional anisotropy** (FA, middle) represents the variability of diffusion in different directions. This value is unit-less, and bounded between 0 and 1 and it is highest in voxels containing a single dense fascicle, such as in the corpus callosum. The **principal diffusion direction** (PDD, right) is the direction of maximal diffusion in each voxel. It is often coded with a mapping of the X, Y and Z components of the direction vector mapped to R, G and B color channels respectively, and scaled by FA. Main white matter structures such as the corpus callosum (red, along the right-to-left x axis) and the corticospinal tract (blue, along the superior-inferior y axis) are easily detected in these maps.

The interpretation of tensor-derived statistics is not straight-forward. MD has been shown to be sensitive to the effects of stroke during its acute phases (Mukherjee, 2005), and loss of nerve fascicles due to Wallerian degeneration and demyelination also results in a decrease in FA (Beaulieu et al., 1996). These changes presumably occur because of the loss of density in brain tissue. The decrease in FA can be traced directly to the loss of myelin structure, which allows for more diffusion of extra-cellular water. But FA also decreases in voxels in which there are crossing fascicles (Pierpaoli and Basser, 1996; Frank, 2001, 2002), rendering changes in FA ambiguous. A recent study estimated that approximately 90% of the white matter contains crossings of multiple fascicles populations (Jeurissen et al., 2013). Therefore, it is unwarranted to infer purely from high FA measures of myelination or in some individuals that these individuals have higher “white matter integrity” (Jones et al., 2013).

In an analogous issue, in voxels with crossing fascicles, the PDD will report the weighted average of the individual fascicles directions, rather than the direction of any one of the fascicles (Rokem et al., 2015). These issues have led to increasing interest in models that represent multiple fascicles within a voxel. Nevertheless, despite its limitations, the statistics derived from the diffusion tensor model provide useful information about tissue microstructure. This model has been so influential that “Diffusion Tensor Imaging” or “DTI” is often used as a synonym for dMRI.

#### Crossing fiber models

One limitation of the diffusion tensor model that was recognized early on (Pierpaoli and Basser, 1996) is that it can only represent a single principal direction of diffusion. Starting with the work of Larry Frank (Frank, 2001, 2002), there have been a series of models that approximate the dMRI signal in each voxel as a mixture combined from the signals associated with different fascicles within each voxel (Tournier et al., 2004; Behrens et al., 2007; Dell’Acqua et al., 2007; Tournier et al., 2007; Rokem et al., 2015)

(see code example: http://arokem.github.io/visual-white-matter/SFM). Common to all these models is that they estimate a fascicle orientation distribution function (fODF), based on the partial volumes of the different fascicles contributing to the mixture of signals. These models are more accurate than the diffusion tensor model in regions of the brain where large populations of nerve fascicles are known to intersect, and also around the optic radiations (Figure 4) (Alexander et al., 2002; Rokem et al., 2015)

**Figure 4:**
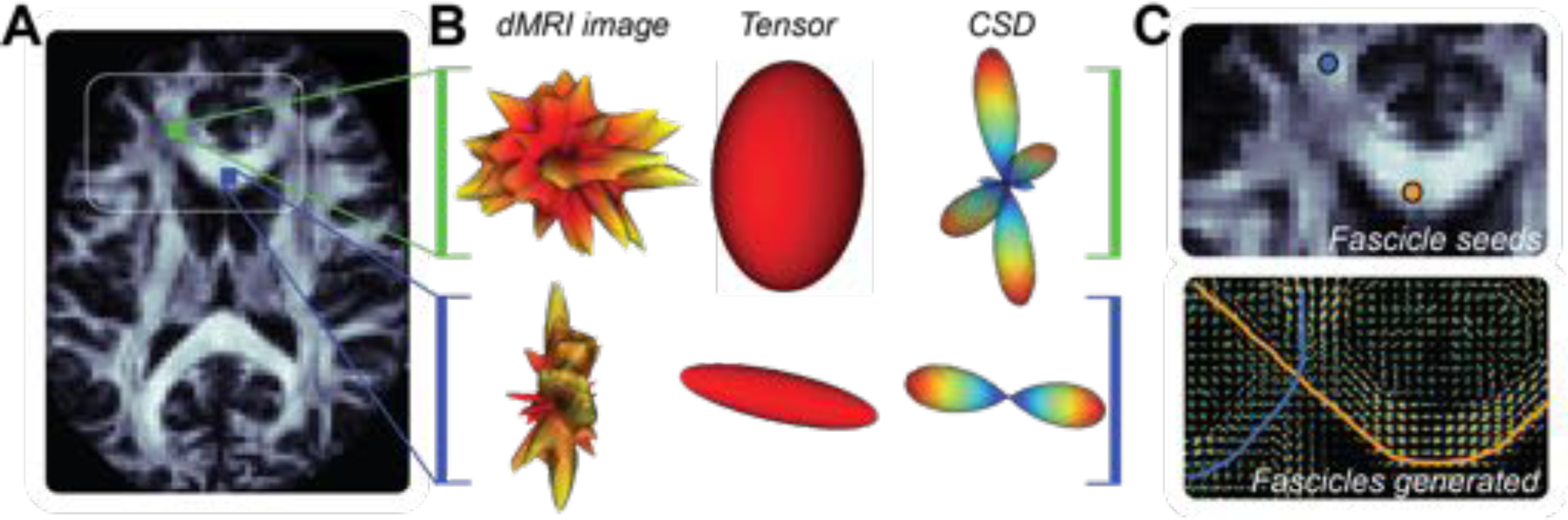
Relation between diffusion MRI signals, voxel models of diffusion and tractographyA. An axial brain slice. Two voxels locations are indicated in blue and green. The white rectangle indicates the location of the images in **C**. **B** Measured diffusion-weighted MR image in the colored locations in A. Left column. Diffusion signal for the green (top) and blue (bottom) locations rendered as 3D surfaces, with the color map indicating the intensity of the diffusion-weighted signal in each direction (red is low signal or high diffusion and yellow-white is high signal, or low diffusion). *Middle column*: The three-dimensional Gaussian distribution of diffusion distance, estimated from the DTM for the signal to the left. The major axis of the ellipsoid indicates the principal diffusion direction estimated by the tensor model, different for the two voxels. *Right column*: Fascicle orientation distribution function (fODF) estimated by a fascicle mixture model: the constrained spherical deconvolution (CSD) model (Tournier et al., 2007) from the signal to its left. CSD indicates several probable directions of fascicles. Colormap indicates likelihood of the presence of fascicles in a direction. **C** *Top panel*: Detail of the region highlighted in **A** (white frame) with example of two seeds randomly placed within the white matter and used to generate the fascicles in the bottom panel. *Bottom panel*: Fits of the CSD model to each voxel and two example tracks (streamlines) crossing at the location of the centrum semiovale.

A second approach to modeling complex configurations of axons in a voxel are offered by so-called “model free” analysis methods, such as q-space imaging (Tuch, 2004) and diffusion spectrum imaging (DSI; (Wedeen et al., 2005)). These analysis methods estimate the distribution of orientations directly from the measurement, using the mathematical relationship between the dMRI measurement and the distribution of diffusion in different directions, and without interposing a model of the effect of individual fascicles and the combination of fascicles on the measurement. These approaches typically require a larger number of measurements in many diffusion directions and diffusion weightings (b-values), demanding a long measurement duration.

### Estimating white matter macrostructure: methods for tracking long-range brain fascicles and connections

Computational tractography refers to a collection of methods designed to make inferences about the macroscopic structure of white matter pathways (Figure 5C; (Jbabdi et al., 2015): these algorithms combine models of the local distribution of neuronal fascicles orientation across multiple voxels, to “track” long-range neuronal pathways. These pathways are generally interpreted as aggregated neuronal projections fascicles, rather than individual axons: brain “highways” that connect distal brain areas, that are millimeters to centimeters apart. Below we describe tractography by dividing it into three major phases of analysis: initiation, propagation, and termination of tracking.

**Figure 5:**
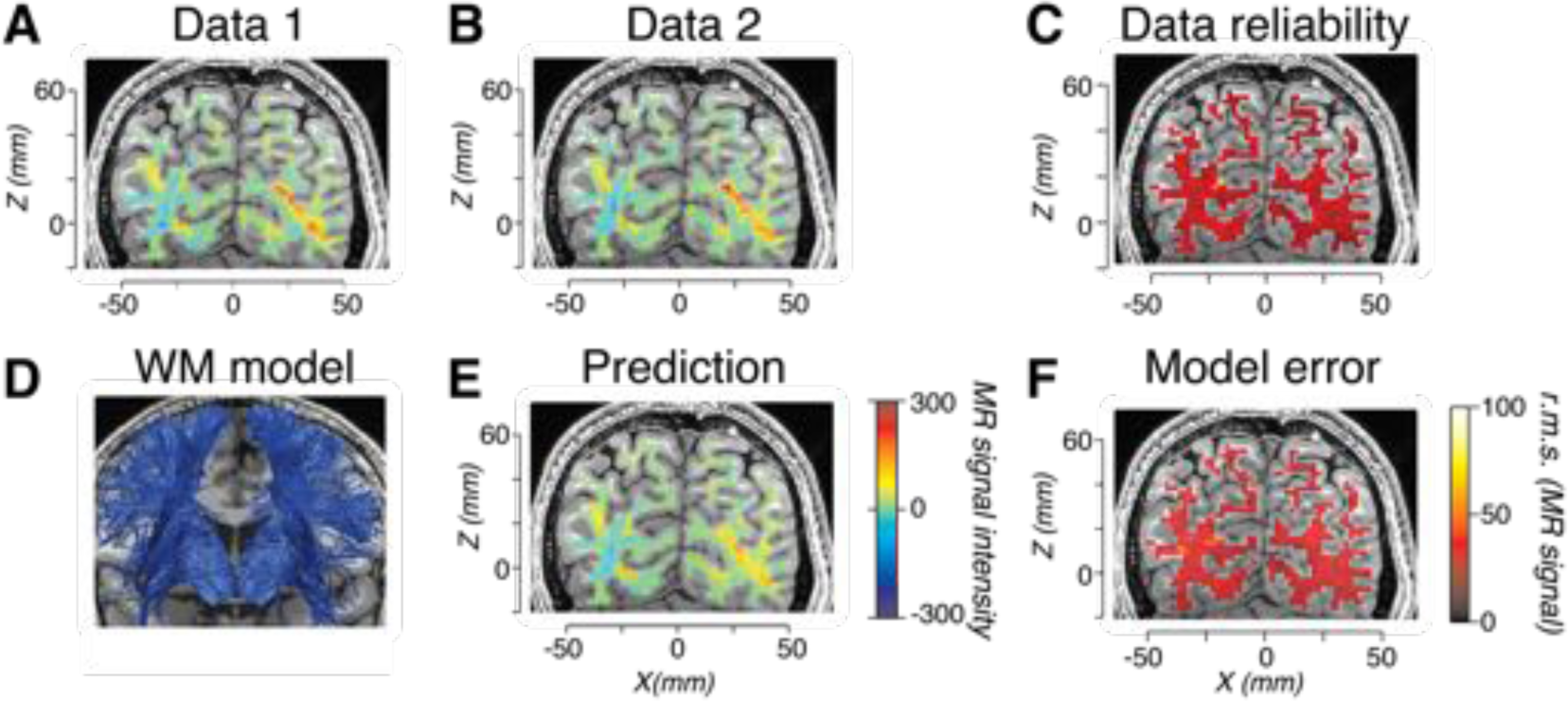
Models of white matter anatomy and tractography evaluation. **A** Measurement of a single diffusion direction shown on a coronal slice of a living human brain. **B**. Repeated measurement of the same direction in **A**, collected during the same scanning session, using the same scanner, sequence, and subject. **C** The reliability of the data, the root mean difference comparing Data 1 and 2 expressed by the color map. **D**. Portion of a full-brain connectome model, comprising part of the corona radiata. Fascicles estimated using Data 1 (**B**) and CSD-based probabilistic tractography. E. Predicted anisotropic diffusion from the model in D, diffusion direction shown is the same as in A and B. Diffusion prediction obtained using the linear fascicle evaluation method, LiFE (Pestilli et al., 2014). **F**. Connectome model accuracy. Root mean-squared error between Data 2 and the model prediction (**E**). Modified from (Pestilli et al., 2014).

#### Track initiation

The process of tractography generally starts by seeding a certain arbitrary number of fascicles within the white matter tissue identified by methods for tissue segmentation (Fischl, 2012). Tracking is generally initiated in many locations within the white matter volume. In several cases, fascicles are seeded within spatially constrained regions of interest. This is the case when the goal is to identify brain connections by tracking between two specific brain areas (Smith et al., 2012).

#### Track propagation

Three major classes of algorithms are used to propagate tracks: deterministic, probabilistic and global. *Deterministic tractography* methods take the fascicles directions estimated in each voxel at face value and draw a streamline by following the principal fascicle directions identified by the voxel-wise model fODF (see above and http://arokem.github.io/visual-white-matter/det_track).

*Probabilistic tractography* methods (Conturo et al., 1999; Mori et al., 1999; Mori and Van Zijl, 2002; Behrens et al., 2003, 2007; Tournier et al., 2012)) accept the voxel-wise estimate of the fODF, but recognize that such estimates may come with errors. Instead of strictly following the directions indicated by the fODF, they consider the fODF to be a probability distribution of possible tracking directions. These methods generate tracks aligned with the principal fiber directions, with higher probability, but they also generate tracks away from the principal directions with nonzero probability (code example: http://arokem.github.io/visual-white-matter/prob_track). *Global tractography* methods build streamlines that conform to specified ‘global’ properties or heuristics (Mangin et al., 2013; Reisert et al., 2011). For example, some algorithms constrain streamlines to be spatially smooth (Aganj et al., 2011).

#### Track termination

Tractography is generally restricted to the white matter volume, so when tracks reach the end of the white matter they are terminated. To determine the extent of the white matter, we can rely on: methods of tissue segmentation based on anatomical measurements (Fischl, 2012) (see http://arokem.github.io/visual-white-matter/det_track for an example) or differences in diffusion properties between different tissue types. For example, because gray matter has low FA, tracking is often terminated when FA drops below a specific threshold (historically FA values of 0.2 or 0.1 have been used as tracking stopping thresholds).

### The Fascicle Detection Problem

Computational fascicle estimated using tractography should be considered as putative models of the structure of neuronal fascicles. Tractography can and does make mistakes and we are far from obtaining gold standard estimates of white matter anatomy and connections using these computational methods. To date, we know that we can validly identify major white matter tracts (Wakana et al., 2007; Yeatman et al., 2012). Yet the fundamental and fine properties of the white matter anatomy are still unknown and matter of debate (for example see (Wedeen et al., 2012a; Catani et al., 2012; Wedeen et al., 2012b)). There is a difficulty in the field in the ability to validate white matter models. Ideally, tractography would be validated routinely in each individual (on a case-by-case basis; Pestilli et al. (2014); Pestilli and Franco (2015)). In the absence of the needed methods several approaches have been proposed to tractography validation.

Post-mortem validation methods have been proposed to show the degree to which tractography can and cannot identify connections found by tracing and histological methods. Unfortunately, such approach cannot contribute to in-vivo understanding of brain function and disease progression. Furthermore, even if multiple brains were to be available for direct inspection with histological and anatomical methods, validation would be difficult as histological validation methods have their own limitations (Goga and Türe, 2015; Simmons and Swanson, 2009; Bastiani and Roebroeck, 2015). Thus, while the ability of most methods to identify major tracts has been validated, it remains difficult to determine whether estimates of tract differences within groups or individuals as measured using dMRI genuinely and accurately correspond to anatomical differences. This has led to interest in more direct methods of validation, such as validation with phantoms of known structure or with simulations (Côté et al., 2013).

Recently, methods that directly estimate the error of a tracking method with respect to the MRI data have been proposed (Daducci et al., 2015; Pestilli et al., 2014; Sherbondy et al., 2008a; Smith et al., 2015). These methods all rely on a forward model approach: the estimated tracts are used to generate a synthetic version of the measured diffusion signal and to compute the tractography model error, by evaluating whether the synthetic and measured data match each other (Figure 5; code example: http://arokem.github.io/visual-white-matter/fascicle-evaluation).

These methods allow evaluation of tractography but not necessarily validation of the anatomy of the final results (Daducci et al., 2016). Relating tractography estimates to brain structures can be conceptualized within the established signal detection theory framework (Green and Swets, 1966). The primary goal of tractography is to detect white matter fascicles and connections. Figure 6 shows the conceptualization of tractography as a signal detection process: every time we track in a brain, there are three potential outcomes of the process; it can identify incorrect fascicles (false alarm), it can miss real fascicles (miss) or it can identify correct white matter fascicles (hits). We call this the *fascicle detection problem* or FDP.

**Figure 6:**
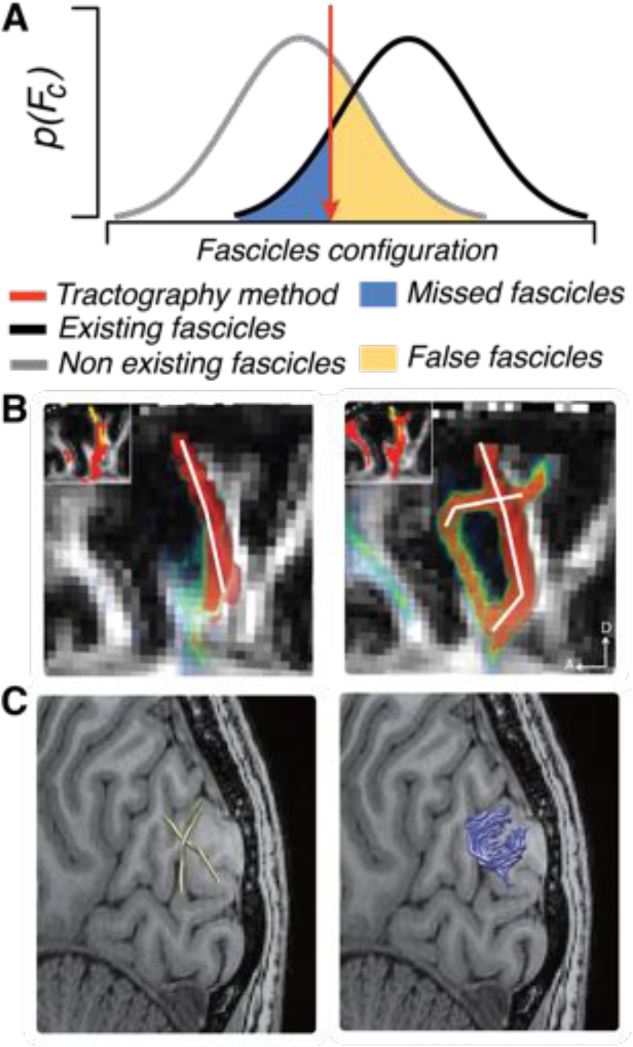
Identifying white matter tract and the signal detection process. **A** The process of tractography described using Signal Detection Theory. The brain contains a certain number of true fascicles (black). The resolution of the diffusion data (either spatial or directional resolution) can provide ambiguous signals indicating fascicles configurations for both existing (black) and non-existing fascicles configurations (gray). Tracking algorithms allow identifying some existing fascicles (right-hand side of the red criterion line), but can miss others (blue highlight), and sometimes can track non-existing fascicles (yellow). **B**. Example of increasing sensitivity for detecting complex fascicles configurations. Left, lower resolution data (1.25 *mm*^3^). Right, higher resolution data (0.7 *mm*^3^). Increasing data resolution improves fascicles detection. This process is akin to changing d’ in the distributions in **A**. **C** Given a fixed data resolution (1.5 mm3 in the example); (Takemura et al., 2016)) changing tractography methods can allow detecting more fascicles, this is akin to a change in criterion in A. In the figure whereas using a conservative turning radius (2 mm) for the streamlines being generated does not allow capturing short-range u-fiber systems (green), using more liberal turning radius (0.5 mm) allows capturing fine details of the anatomy such as short range u-fibers (blue)

Deterministic and probabilistic methods introduced above, use different criteria for the FDP. Deterministic tractography uses a conservative criteria, where it is assumed that each voxel contains only one or few fascicles. The false alarm rate is lower but the miss rate is also higher. The hit rate may also be low, if the fODF used in each voxel does not properly describe the biology of the white matter. Probabilistic tractography uses a more liberal criterion because fascicles can be created using the full extent of the fODF in each voxel. These methods create a higher number of false alarms, but are also more likely to generate hit fascicles (i.e., identify existing fascicles; see also Cote et al. (Côté et al., 2013) for a classification of tractography errors). Global tractography methods lie somewhere between deterministic and probabilistic methods. Their position in the continuum between the very conservative deterministic methods and the very liberal probabilistic methods depends on the details of the global tractography algorithm.

The process of fascicle detection is akin to signal interpretation processes in many other scientific fields where data resolution and signal-to-noise ratio (SNR) limit what can be measured and where methods for data analysis limit how well the signals in the data can be exploited to make inferences. Diffusion data resolution (both angular, the number of diffusion directions measured and spatial resolution, the size of the voxels) and tractography methods (probabilistic vs. deterministic, for example) can change the configuration of fascicles that can be detected. Major white matter tracts can be detected using lower-resolution data. Smaller fascicles might require higher resolution data (Vu et al., 2015). For example, because the the white matter around visual cortex is thin and convoluted, lower resolution dMRI data fails to provide information about connections between the two (dorsal and ventral) banks of the calcarine sulcus. Changing the spatial resolution of the data (e.g., by using a ultra high-field scanner) allows the same tractography methods to find connections between the dorsal and ventral calcarine sulcus (Figure 6B). At the same time, for a given data-set (with its spatial and angular resolution, and SNR) changing tractography methods can change the white matter pathways that are identified. For example, Fig. 6C (left) shows how a “u-fiber” system connecting between human dorsal V3A and V3 can be missed when a conservative curvature parameter is used in probabilistic tractography. Changing the parameters of curvature allows detection of short “u-fibers”, such as the one in Fig. 6C (right). Changing the resolution of the data (number of measured diffusion directions or spatial resolution) is akin to changing d’ (Fig. 6A): increased data resolution improves the sensitivity of the diffusion data, disambiguating signals generated by complex fascicle configurations. Changes in tractography method or parameters may reduce the missed fascicles or the false alarms, but it does not speak to increased sensitivity in detection of white matter pathways. The dilemma is that the best analysis method is likely to depend on the properties of the data as well as the fascicle to be estimated.

A recently-proposed procedure combines the best characteristics of fascicles generated by multiple tractography algorithms (Takemura et al., 2016). The method, called Ensemble Tractography, generates multiple tractography solutions, based on different tractography algorithms. Fascicle evaluation (Pestilli et al., 2014) is then used to arbitrate between the different solutions provided by the different algorithm settings, allowing the method to capitalize on several different levels of sensitivity, while maintaining an accurate representation of the measured signal. Figure 6C shows that this method identifies complex fascicles configurations, easily missed by a single tractography method.

### Comparing tracks and connections across individuals

Once white matter tracts and brain connections have been identified using tractography these estimates have traditionally been analyzed in three ways:

1. Identify major white matter pathways (Catani and Thiebaut de Schotten, 2008; Catani et al., 2002). One typical step after tractography is to cluster these curves together into groups (Garyfallidis et al., 2012; Wassermann et al., 2010) align these curves to each other, either across different individuals, or across hemispheres (Garyfallidis et al., 2015), or to standard anatomical landmarks (Wakana et al., 2007; Yeatman et al., 2012; Yendiki et al., 2011). For example, the optic radiation (Kammen et al., 2015; Sherbondy et al., 2008b) and other connections between thalamus and visual cortex (Ajina et al., 2015; Allen et al., 2015) can systematically be identified in different individuals based on their end-points.
2. Estimate micros-tructural tissue properties (such as FA and MD) within white matter tracts (Yeatman et al., 2012; Yendiki et al., 2011) (see Figure 7A and B).
3. Estimate connectivity between different regions of the cortex (Jbabdi et al., 2015; Rubinov et al., 2009) (See Figure 7C).

**Figure 7:**
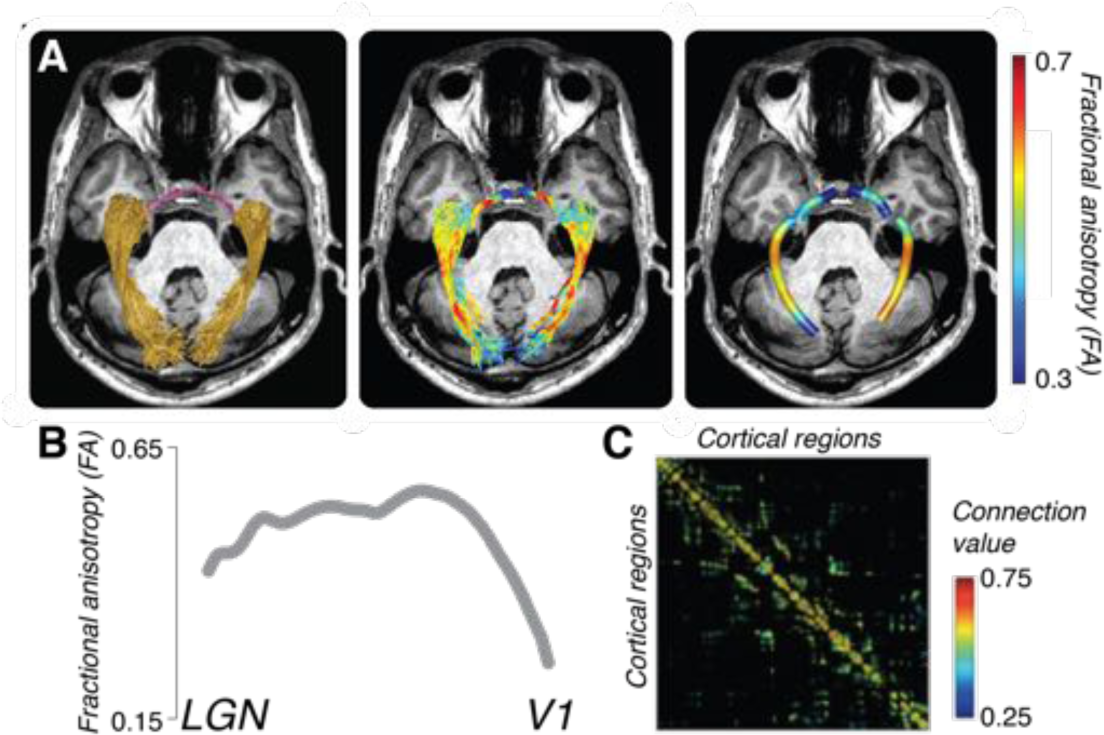
Example applications of tractography to study brain connections and white matter A. Estimates of the optic radiation (OR; left panel gold) and tract (OT; left panel, magenta) from probabilistic tractography (Ogawa et al., 2014). Estimates of fractional anisotropy (FA) projected on top of the anatomy of the OR and OT (right panel). **B** Measurement of FA averaged across the length of the left optic radiation in A. **C**. Matrix of connections between several cortical brain areas. The connection value in each cell of the matrix was derived by normalizing the number of fascicles touching two regions (Sporns, 2013).

All these estimates are combined across individuals and either groups of individuals are compared for statistical significance (for examples by comparing patient groups to control groups) or the estimates are correlated with phenotypic properties and behavior to find correlations between white matter properties and other aspects of human health and behavior of interest. Below we present multiple cases of how such group analyses have been applied to study visual function.

## 2 Tracts and connections across human visual maps

The spatial arrangement of retinal inputs is maintained in visual cortex; signals from nearby locations in the retina project to nearby locations in cortex (Henschen, 1893; Holmes and Lister, 1916; Inouye, 1909; Wandell et al., 2007; Wandell and Winawer, 2011). Vision scientists routinely use functional magnetic resonance imaging (fMRI) to identify these visual field maps (Engel et al., 1994; DeYoe et al., 1994; Sereno et al., 1995; Dumoulin and Wandell, 2008), and more than 20 maps have been identified to date (Smith et al., 1998; Press et al., 2001; Brewer et al., 2005; Larsson and Heeger, 2006; Amano et al., 2009; Arcaro et al., 2009; Silver and Kastner, 2009; Wandell and Winawer, 2011). Recent advances in dMRI and tractography are opening new avenues to study connections between visual field maps through the white matter in living brains. This section introduces what we can learn from relating visual field maps with the endpoints of the white matter pathways.

### Segregation, clustering and hierarchy in visual field maps

The functional organization of the visual field maps in humans and animal models has been identified by convergent knowledge from fMRI, neuropsychological and cytoarchitectonic studies (Wandell et al., 2007; Wandell and Winawer, 2011; Vanduffel et al., 2014; Kolster et al., 2014). We propose three organizational principles for the architecture of the visual field maps to summarize findings in the literature: segregation, clustering and hierarchy. The first principle proposes that visual maps have some degree of functional specialization. This is a soft principle because many areas deal with multiple aspects of the visual stimulus, but some areas respond with more stimulus specificity than others (e.g. motion selectivity in macaque MT and human MT+; (Dubner and Zeki, 1971; Movshon et al., 1985; Watson et al., 1993; Tootell et al., 1995; Huk and Heeger, 2002). There is a classic proposal that functional segregation might extend beyond single visual maps. It has been proposed that visual field maps can be grouped into a dorsal and a ventral stream (Ungerleider and Mishkin, 1982; Ungerleider and Haxby, 1994; Milner and Goodale, 1995). According to this model, the dorsal stream engages in spatial processing and action guidance, whereas the ventral stream engages in analyzing colors and forms. The second principle organizes visual field maps into clusters which share common eccentricity maps (Wandell et al., 2007; Kolster et al., 2014, 2009) and are distinguished by their angular representations. The third principle identifies a hierarchy of visual field maps (Felleman and Van Essen, 1991) based on the input and output layer projections. The first and third principles were defined mostly based on tracer injections, neurophysiological measurements and cytoarchitecture in the macaque brain. The second principle was discovered using functional MRI which has a large field of view and thus can visualize large-scale topological organization of visual field maps.

Functional segregation, clustering and hierarchy are likely to be established by both cortical neuronal sensitivity properties and the connectivity of neurons communicating across different maps. To date, the full anatomical structure of connections of the human visual cortex has not been characterized. This structure is likely to differ between macaque and human brain because the visual field map organization and cortical mantle anatomy differ between the two species in several ways (Vanduffel et al., 2014; Wandell and Winawer, 2011).

Understanding the network of communication pathways in the human brain is fundamental to understanding visual function in health and disease. Building a model of the communication architecture will also help with both prognosis and intervention, this is because vision is a sense that integrates information across many visual features (Huk and Heeger, 2002). For example, skilled reading requires the integration of information across visual space, spatial attention, eye movements, as well as word recognition (Vidyasagar and Pammer, 2010). In fact, there is converging evidence showing the possibility of communication between dorsal and ventral streams (Fang and He, 2005; Grill-Spector et al., 1998, 2000; James et al., 2002; Konen and Kastner, 2008), although the anatomy of white matter pathway communicating these maps clusters was not well understood until very recent reports (see below).

### Relating structure to function: Combining fMRI and diffusion MRI to identify the tracts communicating between maps

There are several examples combining fMRI and dMRI to clarify the relationship between the white matter tracts and visual field maps. Dougherty and colleagues analyzed the relationship between the visual field maps and corpus callosum fascicles (Dougherty et al., 2005). This analysis was extended to measure the properties of this pathway in relation to reading (Dougherty et al., 2007) and blindness (Saenz and Fine, 2010; Bock et al., 2013). Kim and colleagues explored white matter pathways between early visual cortex and category selective regions by combining fMRI, dMRI and fiber tractography (Kim et al., 2006). Greenberg and colleagues identified the visual field maps in intraparietal sulcus using fMRI (Silver and Kastner, 2009) and analyzed the tracts communicating IPS maps and early visual field maps (Greenberg et al., 2012). There are studies attempting to identify the pathway between subcortical nuclei (LGN or Pulvinar) and cortical maps (Ajina et al., 2015; Alvarez et al., 2015; Arcaro et al., 2015).

#### Validating novel tracts

Whereas these approaches provide novel insight on the relationship between white matter pathway and functionally-defined cortical maps, a statistical evaluation can clarify how much the white matter connection between maps are supported by dMRI data increasing the confidence in visual white matter organization identified by dMRI. Pestilli and colleagues developed a method in which the evidence in support of a fascicle of interest is quantified by comparing model accuracy with and without the fascicle (Pestilli et al., 2014). This analysis is used to evaluate the statistical evidence for relatively unstudied pathways, including pathways between visual areas (Gomez et al., 2015; Leong et al., 2016; Takemura et al., 2015).

We used this approach to study the Vertical Occipital Fasciculus (VOF). The VOF, which connects dorsal and ventral visual field maps, was known to 19th century anatomists from postmortem studies (Wernicke, 1881; Sachs, 1892; Déjerine, 1895), but it was widely ignored in the vision literature until recent studies primarily using dMRI (Yeatman et al., 2013; Martino and Garcia-Porrero, 2013; Yeatman et al., 2014b; Takemura et al., 2015; Duan et al., 2015; Weiner et al., 2016). By combining fMRI and dMRI it is possible to visualize the VOF endpoints near the visual field maps (Takemura et al., 2015).

Figure 8C describes the endpoints of the VOF overlaid on the cortical surface together with visual field map boundary identified in Figure 8A. We plotted the position of grey matter nearby the tract endpoints in color maps on cortical surface. The dorsal endpoints of the VOF are near V3A, V3B and neighboring maps (Figure 8C; (Takemura et al., 2015)). The ventral endpoints of the VOF are near hV4 and VO-1. This is important because it sheds light on the nature of communication through the VOF: hV4 and VO-1 are the first full hemifield maps in the ventral stream (Wade et al., 2002; Brewer et al., 2005; Arcaro et al., 2009; Winawer and Witthoft, 2015; Winawer et al., 2010), while V3A and V3B are the first visual field maps to contain a full hemifield representation in the dorsal stream. V3A and V3B also have an independent foveal cluster (Smith et al., 1998; Press et al., 2001) and respond selectively to motion and to disparity (Tootell et al., 1997; Backus et al., 2001; Tsao et al., 2003; Nishida et al., 2003; Ashida et al., 2007; Cottereau et al., 2011; Fischer et al., 2012; Goncalves et al., 2015).

**Figure 8:**
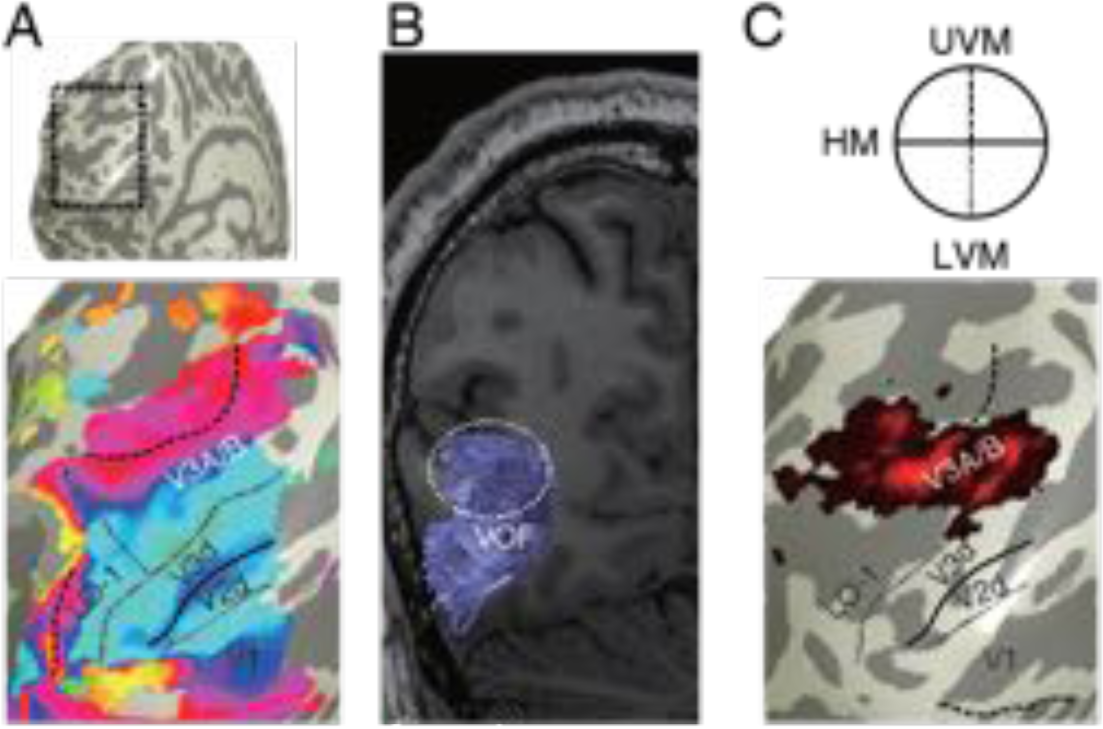
Relation between cortical visual field maps and tract endpoints. **A** Visual field maps identified by fMRI. The surface of dorsal visual cortex was extracted (dotted box in top panel) and enlarged (bottom panel). Colors on the bottom panels describes polar angle representation in population receptive field maps (Dumoulin and Wandell, 2008) red: upper vertical meridian, blue: horizontal, green: lower vertical meridian). The boundaries between visual field maps are defined as polar angle reversals in population receptive field maps. **B** The VOF identified in an identical subject using dMRI and fiber tractography. Reproduced from (Takemura et al., 2015) with permission. **C** Overlay between visual field maps and VOF projections. Color maps indicate voxels near the VOF endpoints in this hemisphere, and the color scale indicates the density of fascicle endpoints. The borders of visual field maps (solid and dotted lines; upper panel depicts the representation of horizontal and vertical meridians) were adopted from fMRI-based mapping (**A**).

We note that there are some limitations to associating “tract endpoints” in tractography and cortical projections in gray matter. In practice, tractography is usually terminated at the boundary between gray and white matter, because standard dMRI measurements fall under the resolution required to accurately delineate the complex tissue organization within gray matter. Therefore, there is uncertainty in relating tract endpoints to the cortical surface. This uncertainty is compounded by the dispersion of axons in gray matter, and by U-fiber system in superficial white matter which impedes accurate estimation of tract endpoints (Reveley et al., 2015). Thus, the analysis will give us the information on the spatial proximity between tract endpoint and cortical maps, but does not provide us a definitive estimate on the cortical projection of the tract. We may expect the improvement of the quality of this analysis by utilizing the modelling of complex fiber organization from diffusion MRI data, improving data resolution (Sotiropoulos et al., 2013) or improved tractography methods (Reisert et al., 2011; Takemura et al., 2016; Wandell, 2016).

## 3 Relationship between the visual white-matter and visual impairment

One of the motivations for studying the effects of blindness on white matter, is that our extensive understanding of the anatomical and functional structure of the visual system, both in visually normal and in visually deprived animals, provides a good model system in which to evaluate current methods for measuring white matter structure in vivo. Indeed since the classic studies of Hubel and Wiesel in the 1960s (Wiesel and Hubel, 1963, 1965b, a; Wiesel et al., 1963), visual deprivation has been an important model for examining the effects of early experience on brain development and function. In this section, we focus on the effects of visual loss, due to peripheral blindness (such as retinal disease), on white matter pathways. In addition, we discuss how in vivo measurements of white matter structure can provide insight into pathways that mediate blindsight after striate lesions. Blindsight provides a particularly elegant example of determining which of multiple candidate pathways are likely to mediate residual behavioral performance based on in vivo measurements of white matter structure.

### White matter changes due to peripheral blindness

Visual deprivation resulting from loss of input from the eye provides an excellent model for understanding the links between structure and function, as the brain itself remains largely intact. A wide variety of studies in both animals and humans have demonstrated large scale functional (Lewis and Fine, 2011), neuroanatomical (Bock and Fine, 2014; Movshon and Van Sluyters, 1981), neurochemical (Coullon et al., 2015; Weaver et al., 2013) and even vascular (De Volder et al., 1997; Uhl et al., 1993) changes within occipital cortex as a result of early blindness. Given this extensive animal and human literature, peripheral blindness provides a particularly good model system to examine the reliability and sensitivity of current noninvasive methods for assessing the effects of experience on white matter pathways. Here, we review the white matter changes associated with visual deprivation, paying particular attention to the links between white matter microstructure and alterations of functional responses due to visual loss.

### The optic tract and optic radiations

Early or congenital blindness results in severe degradation of the optic tract (between the eye and the lateral geniculate nucleus) and the optic radiations (between the lateral geniculate nucleus and V1). Not only are these pathways noticeably reduced in volume, but there is also decreased longitudinal and increased radial diffusivity within these tracts (Noppeney et al., 2005; Ptito et al., 2008; Reislev et al., 2015; Shu et al., 2009) suggesting degradation of white matter microstructure within the remaining tract. These effects seem to be particularly pronounced in anophthalmic individuals, suggesting that retinal signals play a role in the development of these tracts (Bridge et al., 2009).

Reductions in fractional anisotropy (FA) within the optic tract and optic radiations are also found in individuals who become blind in adolescence or adulthood (Dietrich et al., 2015; Shimony et al., 2006; Wang et al., 2013). There is some suggestion that the microstructure of these tracts may be more heavily degraded in individuals with acquired blindness than in congenitally blind individuals, though in some part these results may be due to the etiologies of later blindness - for example glaucoma is capable of causing physical damage to the optic tract.

### Callosal connectivity

Despite numerous studies, the effects of early visual deprivation on the posterior callosal fibers within the splenium (connecting the occipital lobes) are not yet fully understood. Several studies (Leporé et al., 2010; Ptito et al., 2008; Tomaiuolo et al., 2014) have reported that the posterior part of the corpus callosum is reduced in volume. However, Bock et al. (2013) failed to find any indication of reduced volume in the portion of the splenium containing fibers connecting V1/V2 in either early blind or anophthalmic individuals (Bock et al., 2013). In the Bock et al. study, the only evidence of callosal atrophy due to visual loss was a slight reduction in FA within the splenium in anophthalmic individuals. Given the magnitude of the effect found by Tomaiuolo et al., the Bock study may have been underpowered, though there was no indication of a non-significant difference between groups. An alternative possibility is that the significant methodological differences across the studies may explain the discrepancy in results. Most of the studies listed above registered individual data to a template, with some studies then anatomically segmenting the callosum into sub-regions of various sizes. In contrast, Bock et al. defined the splenium in a midline sagittal slice using fibers tracked from a surface based ROI, in which V1/V2 was identified based on cortical folding for each subject – resulting in a much more restrictive definition of the splenium. Indeed, the splenial region of interest in the Bock et al. (2013) study was roughly a third of the area defined by Tomaiuolo et al. (Tomaiuolo et al., 2014). Thus, one possible explanation for these differing findings is that the callosal fibers that connect V1/V2 are less affected by visual deprivation than those representing higher-level cortical areas.

As well as examining the volume of the splenium, Bock et al. (2013) also showed that the topographic organization of occipital fibers within the splenium is maintained in early blindness. Even within anophthalmic individuals, it was possible to observe a ventralâĂŠSdorsal mapping within the splenium with fibers from ventral V1-V3 (representing the upper visual field) projecting to the inferior-anterior corner of the splenium and fibers from dorsal V1-V3 (representing the lower visual field) projecting to the superior-posterior end. Similarly, it was possible to observe an eccentricity gradient of projections from foveal-to peripheral V1 subregions running in the anterior-superior to posterior-inferior direction, orthogonal to the dorsal-ventral mapping(Bock et al., 2013).

### Cortico-cortical connections

Over the last two decades a considerable number of studies have observed functional cross-modal plasticity (novel or augmented responses to auditory or tactile stimuli within occipital cortex) as a result of congenital blindness. Specifically, occipital regions have been shown to demonstrate functional responses to a variety of auditory (Röder et al., 1999; Lessard et al., 1998; Voss et al., 2004; Gougoux et al., 2004; Jiang et al., 2014), language (Sadato et al., 1996; Cohen et al., 1997; Röder et al., 2002; Burton et al., 2003; Bedny et al., 2011; Watkins et al., 2012) and tactile (Alary et al., 2009; Goldreich and Kanics, 2003; Van Boven et al., 2000) stimuli. One of the more attractive explanations for these cross-modal responses, given the animal literature (Innocenti and Price, 2005; Restrepo et al., 2003; Webster et al., 1991), is that they might be mediated by altered white matter connectivity – for example a reduction in experience-dependent pruning. As a result, when methods for measuring white matter tracts in vivo became available there was immediate interest in looking for evidence of novel and/or enhanced connections within early blind individuals. However, despite the massive changes in functional responses within occipital cortex that are observed as a result of blindness, cortico-cortico white matter changes as a result of early blindness are not particularly dramatic, with little or no evidence for enhanced connectivity as predicted by the ‘reduced pruning’ hypothesis.

Indeed, reduced fractional anisotropy is consistently found within white matter connecting the occipital and temporal lobes (Bridge et al., 2009; Ptito et al., 2008; Reislev et al., 2015; Shu et al., 2009). This finding is somewhat puzzling given that several studies have shown that language stimuli activate both the lateral occipital cortex and fusiform regions in congenitally blind individuals (Bedny et al., 2011; Watkins et al., 2012). Similarly, no increase in FA has been detected in the pathways between occipital and superior frontal cortex, a network that has shown to be activated by language in congenitally blind populations (Bedny et al., 2011; Watkins et al., 2012) (Bedny et al., 2011; Watkins et al., 2012). Indeed two studies have found evidence of deterioration (e.g. reduced FA) within the inferior fronto-occipital fasciculus (Reislev et al., 2015; Shu et al., 2009).

It is not clear what causes the lack of consistency in the studies described above. One likely factor is between-subject variability. Estimates of individual differences in tracts suggest that between-group differences require surprisingly large subject numbers even for relatively large and consistently located tracts. For example, within the inferior and superior longitudinal fasciculus, to detect a 5% difference in fractional anisotropy at 0.9 power would require 12-19 subjects/group in a between-subject design (Veenith et al., 2013). Because of the difficulty of recruiting blind subjects with relatively homogenous visual histories, many of the studies cited above used moderate numbers of subjects.

One of the more puzzling findings described above is the consistent finding of reduced white connectivity between occipital and temporal lobes and between occipital and superior frontal lobes, given that occipital cortex shows enhanced responses to language as a result of blindness. One possibility is that measured connectivity provides a misleading picture of actual connectivity between these regions. One potential confound is the effect of crossing fibers. If blindness leads to a loss of pruning that extends beyond these major tracts then the resulting increase in the prevalence of crossing fibers within other networks might reduce measurable FA within occipito-temporal and occipito-frontal tracts.

Alternatively, it is possible that the enhanced language responses found in occipital cortex genuinely co-exist with reduced connectivity with temporal cortex. For example, we have recently suggested that an alternative perspective in understanding cortical plasticity as a result of early blindness is to consider that occipital cortex may compete rather than collaborate with non-deprived regions of cortex for functional role (Bock and Fine, 2014). According to such a model, lateral occipital cortex may compete with temporal areas, with language tasks assigned to the two regions in such a way as to maximize the decoupling between the areas. In the context of such a framework, reductions in anatomical connectivity between the two areas are less surprising.

### Early vs. late blindness

Reductions in fractional anisotropy (FA) within the optic tract and optic radiations are also found in individuals who become blind in adolescence or adulthood (Dietrich et al., 2015; Shimony et al., 2006; Wang et al., 2013), suggesting that visual input is necessary for maintenance as well as development of these tracts. Indeed, there is some suggestion that the microstructure of these tracts may be more heavily degraded in individuals with acquired blindness than in congenitally blind individuals, though in some part these results may be due to the etiologies of later blindness - for example glaucoma is capable of causing physical damage to the optic tract.

The effects of early vs. late blindness on cortico-cortico white matter connections remains unclear. One study has found that reductions in FA may be more widespread in late acquired blindness compared to congenital blindness, with late blind individuals showing reduced FA in the corpus callosum, anterior thalamic radiations, and frontal and parietal white matter regions (Wang et al., 2013). Such a result is quite surprising, though it is possible that the cross-modal plasticity that occurs in early blind individuals might prevent deterioration within these tracts, and this ‘protective’ effect does not occur in late blind individuals. However this difference between late and early blind subject groups was not replicated by Reislev et al. (Reislev et al., 2015) who noted similar losses in FA for early and late blind subjects within both the inferior longitudinal fasciculus and the inferior fronto-occipital fasciculus using a tract-based approach. (This group did find that significant increases in radial diffusivity within late blind subjects relative to sighted control subjects, but did not directly compare radial diffusivity between late and early blind subject groups.)

### White matter changes due to cortical blindness

Damage to the primary visual cortex (V1) causes the loss of vision in the contralateral visual field (homonymous hemianopia), which in turn leads to degeneration of the geniculo-striate tract. Despite these lesions and a lack of conscious vision, many people can still correctly detect the presence/absence of a stimulus (Poppel et al., 1973), as well as discriminate between variations in stimulus color and motion, (Ffytche et al., 1996; Morland et al., 1999) despite a lack of conscious awareness of the stimulus – a condition known as ‘blindsight’ (Poppel et al., 1973; Weiskrantz et al., 1974). Given the absence of the major geniculo-striate projection, any residual visual information is likely to be conveyed by an intact projection from subcortical visual regions to undamaged visual cortex.

While several individual case studies have examined the white matter pathways underlying residual vision (Bridge et al., 2010, 2008; Leh et al., 2006; Tamietto et al., 2012), a critical test of the functional relevance of white matter microstructure was recently carried out by comparing individuals with V1 lesions who did and did not demonstrate residual ‘blindsight’ vision. Ajina et al. (2015) examined white matter microstructure within three potential pathways that might subserve residual vision in hemianopia patients. They showed that two potential pathways (superior colliculus to hMT+ and the interhemispheric connections between hMT+) did not differ in microstructure between those with and without blindsight. In contrast, the pathway between LGN and hMT+ in the damaged side differed significantly across individuals with and without blindsight. Patients with blindsight had comparable FA and MD in damaged and intact hemispheres. Patients without blindsight showed a loss of structural integrity within this tract in the damaged hemisphere (Ajina et al., 2015). These results provide an elegant example of how white matter microstructure measurements can be correlated with visual function in order to infer structure-function relationships.

### Summary

Within the subcortical pathways, our review of the literature suggests that in vivo measurements of white matter microstructure can provide sensitive and reliable measurements of the effects of experience on white matter. Studies consistently find the expected deterioration within both the optic tract and the optic radiations in the case of peripheral blindness. Similarly, analysis of subcortical pathways in blindsight have shown consistent results across several cases, and are interpretable based on the functional and pre-existing neuroanatomical literature.

In contrast, it has proved surprisingly difficult to obtain consistent results for measurements of cortico-cortico connectivity across different studies. These discrepancies in the literature are somewhat surprising given the gross differences in visual experience that occurs in early and even late blind individuals. One possible explanation is that, as described above, most studies (though with a few exceptions, e.g. Wang et al. (2013)) have tended to have relatively low statistical power. Larger sample sizes (perhaps through a multi-center study) might better reveal more subtle anatomical differences in connectivity as a result of early blindness. A second factor may be methodological differences across studies – early blind subjects are anatomically distinct within occipital cortex – for example occipital cortex shows less folding in animal models (Dehay et al., 1996), which may affect estimates of white matter tracts that are based on methods involving normalization to a canonical template. Comparing results based on tracts identified based on normalization to anatomical templates to those obtained using tracts identified within individual anatomies may provide some insight into how these very different approaches compare in terms of sensitivity and reliability.

It has also proved surprisingly difficult to interpret alterations in white matter cortico-cortico connectivity in the context of the functional literature – occipito-temporal and occipito-frontal white matter connections have consistently been shown to be weaker in early blind subjects, despite the apparent recruitment of occipital cortex for language. This discrepancy between estimates of white matter connectivity and functional role within cortico-cortical tracts makes it clear that drawing direct conclusions from white matter microstructure to functional role is still fraught with difficulty. This is not a reason to abandon the enterprise, but rather provides a critical challenge. To return to the argument with which we began this chapter – the effects of visual deprivation provide an excellent model system for testing how well we understand the measurement and interpretation of in vivo measurements of white matter infrastructure.

## 4 Properties of white matter circuits involved in face perception

In the visual system, neural signals are transmitted through some of the most prominent longrange fiber tracts in the brain: the optic radiations splay out from the thalamus to carry visual signals to primary visual cortex; the forceps major, u-shaped fibers that traverse the splenium of the corpus callosum, innervate the occipital lobes allowing them to integrate neural signals; the inferior longitudinal fasciculus (ILF) is the primary occipito-temporal associative tract (Crosby, 1962; Gloor, 1997) that propagates signals through ventral visual cortex between primary visual cortex and the anterior temporal lobe (see Figure 9a); and, finally, the inferior fronto-occipital fasciculus (IFOF) begins in the occipital cortex, continues medially through the temporal cortex dorsal to the uncinate fasciculus, and terminates in the inferior frontal and dorsolateral frontal cortex (see Figure 9a; Catani et al. (2002). Increasingly, diffusion neuroimaging studies are providing evidence indicating the importance of these white matter tracts for visual behavior. Here, we present evidence from converging streams of our research demonstrating that the structural properties of the ILF and IFOF are critically important for intact face perception.

**Figure 9:**
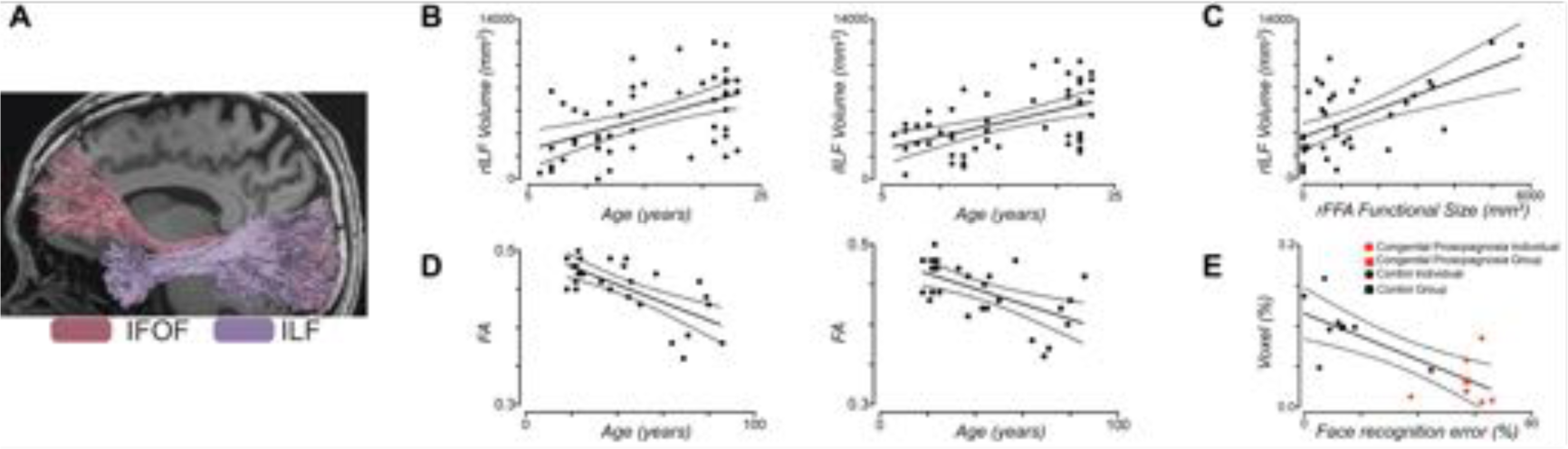
The Role of the ILF and IFOF in face perception. **A** Representative example of fibers extracted from the inferior longitudinal fasciculus (ILF), which is the primary occipito-temporal associative tract (Crosby, 1962; Gloor, 1997) that propagates signals through ventral visual cortex between primary visual cortex and the anterior temporal lobe; and, from the finally, the inferior fronto-occipital fasciculus (IFOF), which begins in the occipital cortex, continues medially through the temporal cortex dorsal to the uncinate fasciculus, and terminates in the inferior frontal and dorsolateral frontal cortex (Catani et al., 2002). **B** In typically developing individuals, there is a systematic age-related increase in the macro-structural properties of both the right and left ILF, which is measured here by the volume of the tract in cubic millimeters, across the first two decades of life. **C** We observed a joint structure-function correspondence between the size of the individually defined functional right fusiform face area and the volume of the right ILF, which held even when age was accounted for (Scherf et al., 2014). **D** In typically developing adults, there is a systematic age-related decline in the micro-structural properties of both the right and left IFOF, as measured by the mean FA, indicating that it is disproportionately vulnerable compared to the ILF during the aging process. **E** Individuals who have a lifetime history of face blindness, an inability to recognize faces in spite of normal intelligence and sensory vision, have systematically smaller volume visual fiber tracts, particularly in the right ILF, as depicted here, compared to age-matched control participants. Together, these findings provide converging evidence that these two major tracts, the IFOF and the ILF, carry signals important for face perception.

Face perception is a complex suite of visual behaviors that is subserved by a distributed network of neural regions, many of which are structurally connected by the ILF and IFOF. For example, the “core” or posterior regions include the occipital face area (OFA), the fusiform face area (FFA), and the posterior superior temporal sulcus (pSTS) and the “extended” areas include the anterior temporal pole (ATP), amygdala, and ventro-medial prefrontal cortex (vmPFC) (Gobbini and Haxby, 2007). Recently, functional neuroimaging studies have provided supporting, albeit indirect, evidence of rapid interactions between these posterior core regions and the extended anterior regions (e.g., anterior temporal lobe and amygdala) that implicate the involvement of long-range association fiber tracts that connect these regions, including the ILF and the IFOF (Bar et al., 2006; Gschwind et al., 2012; Rudrauf et al., 2008; Song et al., 2015).

Finally, damage to either of these pathways disrupts face processing (Catani et al., 2003; Catani and Thiebaut de Schotten, 2008; Fox et al., 2008; Philippi et al., 2009; Thomas et al., 2009), suggesting that these white matter tracts serve as a critical component of the neural system necessary for face processing. In what follows, we provide evidence showing that age-related changes in the structural properties of these tracts in early development is associated with emergent properties of the functional neural network supporting face processing. We also review evidence that age-related decreases in the structural properties of these tracts in aging adults exist and are associated with decrements in face processing behavior. Finally, we review data showing that relative deficits in the structural properties of these long-range fiber tracts potentially explain the causal nature of face blindness in individuals with congenital prosopagnosia. Together these findings converge to indicate that the structural properties of white matter tracts, and the ILF and IFOF in particular, are necessary for skilled face perception.

Across all of these studies, we use a common methodological approach. We acquire diffusion images, analyze them using a tensor model, and perform deterministic tractography using the Fiber Assignment of Continuous Tracking (FACT) algorithm and brute-force fiber reconstruction approach (Mori et al., 1999; Xue et al., 1999) with fairly standard parameters. To extract the tracts of interest, we use a multiple region of interest (ROI) approach that is very similar to Wakana et al (Wakana et al., 2007). From each tract, we extract the volume (the number of voxels through which the fibers pass multiplied by the volume of the voxel) as well as the mean fractional anisotropy (FA), mean diffusivity (MD), axial diffusivity (AD), and radial diffusivity (RD) values across these voxels.

### Age-related changes in the ILF and IFOF with functional neural change

The central question guiding this work was whether developmental differences in the structural properties of the ILF and IFOF, defined independently using anatomical ROIs, are related to developmental differences in the characteristics of the functional face-processing regions connected by these tracts (Scherf et al., 2014). Across participants whose ages covered a substantial range (ages 6-23 years), we evaluated differences in (1) the macro- and microstructural properties of the fasciculi, and in (2) the functional profile (size, location, and magnitude of selectivity) of the face- and place-selective regions that are distributed along the trajectory of the pathways of interest using functional magnetic resonance imaging (fMRI). First, we found that all tracts, with the exception of the left IFOF, exhibited age-related improvements in their microstructural properties, evincing a significant decrease in mean and radial, but not axial, diffusivity from childhood to early adulthood (see Figure 9b). This result is consistent with the idea that the increasingly restricted diffusion perpendicular to the axons in these tracts reflects continued myelination from childhood through early adulthood. The left IFOF exhibited stable levels of microstructural properties across the age range, indicating that it may be a very early developing fiber tract. At the macro-structural level, only the ILF exhibited an age-related change in volume, which was evident in both hemispheres. The combined increases in micro-structural properties and the volume of the right ILF suggest that it is becoming increasingly myelinated and/or more densely packed with axons with age (Beaulieu, 2002; Lebel and Beaulieu, 2011; Song et al., 2005).

Having identified the structural changes that potentially contribute to circuit organization, we explored, concomitant differences in the functional profile of face-related cortical regions, and then examined the joint structure-function correspondences. Across the full age range, individuals with larger right FFA volumes also exhibited larger right ILF volumes (see Figure 9c). Neither the right OFA nor the right PPA exhibited this structure-function relation with the right ILF. This structurefunction relation between the right FFA and right ILF was also present in just the children and adolescents (aged 6-15 years). However, once age was accounted for in the same model as size of the right FFA in the children and adolescents, the age effects on the volume of the ILF swamped all the significant variation, even though the size of the right FFA increased significantly across this age range. One interpretation of these findings is that the neural activity generated by larger functional regions may require and/or influence the development of larger fiber tracts (via increasing myelination of existing axons and/or more densely packed axons) to support the transmission of neural signals emanating from such regions. It may also be that increasing the integrity of the structural architecture of fiber tracts increases the propagation of neural signal throughout the circuit, thereby enhancing the functional characteristics of the nodes within the circuit (and vice versa). In other words, structural refinements of white matter tracts may precede, and even be necessary for, functional specialization of the circuit to emerge. In sum, this work uncovers the key contributions of the ILF and IFOF and their relationship to functional selectivity in the developing circuitry that mediates face perception. Aging related decrease in structural properties of IFOF with face perception deficits While the findings above focus on understanding how the structural properties of the ILF and IFOF are critical for building the complex face processing network during early development, it was also important to understand whether disruptions in these white matter tracts might be associated with, or even responsible for, age-related declines in face processing that are well reported (e.g., Salthouse (2004)) and are not solely a function of memory or learning changes (Boutet and Faubert, 2006). We scanned 28 individuals aged 18-86 years using a diffusion tensor imaging protocol (Thomas et al., 2008). We also tested them in face and car perceptual discrimination tasks. We observed that the right IFOF was the only tract that decreased in volume (as measured by percent of fibers and voxels through which the fibers pass) as a function of age. In contrast, the bilateral IFOF and the left, but not right, ILF exhibited age-related decreases in FA (see Figure 9d). To summarize, it is the IFOF in the right hemisphere that shows particular age-related vulnerability, although there is a tendency for the tracts in the left hemisphere to show some reduction in microstructural properties, as revealed in FA values, as well.

On the discrimination tasks, participants performed more poorly on the difficult trials, especially in the face compared to the car condition. Of relevance though is that the older individuals, the 60and 80-year-olds, made significantly more errors on faces than they did on cars, and performance was almost at chance in the difficult condition. Given that there were age-related declines in the micro- and macro-structural properties of the IFOF as well as in face perception behavior, our final question was whether there was an association between these tract and behavioral deficits. To address this question, we examined correlations between behavioral performance and the normalized percentage of fibers, normalized percentage of voxels and average FA values in the ILF and IFOF in each hemisphere. We observed two main findings: a. during the easy face trials, participants with greater percentage of fibers in the right IFOF, exhibited better performance in these easy discriminations and b. during the difficult face trials, participants with higher FA values and larger volume right IFOF exhibited better performance on the difficult discriminations. Taken together, these findings indicate a clear association between the ability to discriminate between faces and the macro- and micro-structural properties of the IFOF in the right hemisphere.

### Disruptions in ILF and IFOF may explain face blindness

Finally, given the findings of an association between face processing behavior and the structural properties of the IFOF in typically developing adults, we investigated whether congenital prosopagnosia, a condition that is characterized by an impairment in the ability to recognize faces despite normal sensory vision and intelligence, might arise from a disruption to either and/or both the ILF and IFOF (Thomas et al., 2009). This hypothesis emerged following empirical findings that the core functional neural regions, including the FFA, appeared to produce normal neural signals in many congenital prosopagnosic individuals (Avidan et al., 2014, 2005; Thomas et al., 2009). To test this hypothesis, we scanned 6 adults with congenital prosopagnosia, all of whom evinced normal BOLD activation in the core face regions (Avidan et al., 2005), and 17 age- and gender-matched control adults. We also measured participants’ face recognition skills. Relative to the controls, the congenital prosopagnosia (CP) group showed a marked reduction in both the macro- and micro-structural properties of the ILF and IFOF bilaterally. We then carried out a stepwise regression analysis with the connectivity and behavioral measures. Across participants, individuals with the worse face recognition behavior had the lowest FA and smallest volume right ILF (see Figure 10e). Similarly, poor face recognition behavior was also related to smaller volume of the right IFOF, as well. In summary, our study reveals that the characteristic behavioral profile of congenital prosopagnosia may be ascribed to a disruption in structural connectivity in some portion of the ILF and, perhaps to a lesser extent, the IFOF (for related findings, see Gomez et al. (2015); Song et al. (2015)).

### Summary

We have presented converging evidence that face recognition is contingent upon efficient communication across disparate nodes of a widely distributed network, which are connected by long range fiber tracts. Specifically, we have shown that two major tracts, the IFOF and the ILF, carry signals important for face perception. Studies in children and older individuals, as well as investigations with individuals who are face blind all attest to the required integrity of the tracts for normal face recognition behavior. Early in development the ILF undergoes a particularly long trajectory in which both the micro- and macro-structural properties change in ways indicative of increasing myelination. Of relevance for the integrity of face perception behavior, there is a highly selective and tight correspondence between age-related growth in one of the pre-eminent functional nodes of the face-processing neural network, the right FFA, and these age-related improvements in the structural properties of the ILF. These findings reflect the dynamic and intimate nature of the relation between brain structure and function, particularly with respect to role of white matter tracts, in setting up neural networks that support complex behaviors like face perception. Our work with congenital prosopagnosic participants suggests that face-processing behavior will not develop normally when this developmental process is disrupted. Future work will need to identify when, developmentally, white matter disruptions are present and interfere with face perception. Finally, our work in aging adults indicates that the IFOF, is particularly vulnerable with age, which can lead to difficulties with face recognition behavior. However, Grossi and colleagues (Grossi et al., 2014) reported that an adult with progressive prosopagnosia, a gradual and selective inability to recognize and identify faces of familiar people, had markedly reduced volume ILF in the right, but not left hemisphere, whereas the bilateral IFOF tracts were preserved. This suggests that there may be some conditions in adulthood in which the ILF is also vulnerable. Together, these findings converge on the claim that the structural properties of white matter circuits are, indeed, necessary for face perception.

## 5 Conclusions, remarks and future directions

The goal of the present article is to survey the range of findings about the specific importance of studying the white matter of the visual system. We focused on findings in human brains, that used the only currently available method to study the white matter in vivo in humans: diffusion MRI (dMRI). In the time since its invention in the 1990’s, dMRI methods have evolved, and evidence about the importance of the white matter has accumulated (Fields, 2008b). These findings are modern, but they support a point of view about the nervous system that has existed for a long time. Connectionism, the theory that brain function arises from its connectivity structure goes back to classical work of the 19th century neurologists (Deacon, 1989). Over the years, interest in disconnection syndromes waxed and waned, as more holistic views of the brain (Lashley, 1963) and more localizationist views of the brain prevailed. As a consequence, systems neuroscience has traditionally focused on understanding the response properties of individual neurons and cortical regions (Fields, 2013, 2004). Only a few studies were devoted to understanding the relation of the white matter to cognitive function. Similar to electrical cables, it was believed that the connections in the brain were either intact and functioning, or disconnected. However, the pendulum started swinging back dramatically towards connectionism already in the 1960’s (Geschwind, 1965), and over the years, there has been increasing appreciation for the importance of brain networks in cognitive function, culminating in our current era of connectomics (Sporns et al., 2005). In addition to an increasing understanding of the importance of connectivity, views that emphasize the role of tissue properties not related to neuronal firing in neural computation have more recently also evolved and come to the fore (Bullock et al., 2005; Fields, 2008b). For example, the role of glial cells in modulating synaptic transmission and neurotransmitter metabolism, as well as the role of the white matter in metabolism and neural hemodynamic coupling (Robel and Sontheimer, 2016). Today, we know much more about the white matter than ever before and we understand that the properties of the tissue affect how communication between distal brain areas is implemented. Diffusion MRI enables inferences about the properties of the white matter tissue in vivo, which in turn enables inferences about the connection between behavior and biology.

As we have seen in the examples presented above, dMRI is a useful method to study the biological basis of normal visual perception, to glean understanding about the connectivity that underlies the organization of the visual cortex, about the breakdowns in connectivity that occur in different brain disorders, and about the plasticity that arises in the system in response to visual deprivation. But while the findings reviewed above demonstrate the importance of the white matter in our understanding of the visual system and the biological basis of visual perception, they also serve as a demonstration of the unique capacity of vision science to study a diverse set of phenomena across multiple levels of description. This is largely an outcome of the detailed understanding of different parts of the visual system, and attributable to the powerful quantitative methods that have developed in the vision sciences to study the relationship between biology and perception. For these reasons the visual system has historically proven to be a good testing ground for new methodologies. Another recurring theme of the examples presented above is the importance of convergent evidence in cognitive neuroscience (Ochsner and Kosslyn, 1999). Studies of the human visual white matter exemplify this: evidence from behavior, from functional MRI, from developmental psychology, and from physiology converge to grant us a unique view about the importance of specific pathways for perception (see sections 2 and 4). For example, the response properties of different brain regions are heavily influenced by the synaptic inputs of long range connections: one might grow to better understand the response properties of dorsal visual areas, when their connection through the white matter with ventral visual areas is known (see section 2). Similarly, the role of different functional regions in the face-perception network becomes clearer when the relationship between the size of these regions, the size of the ILF and the co-development of these anatomical divisions is demonstrated (see section 4). The study of individuals with perceptual deficits or with visual deprivation demonstrates the specific importance of particular connections, and delineates the lifelong trajectory of the role of these trajectories, also revealing possibilities and limitations of brain plasticity (see sections 3). The importance of converging evidence underscores the manner in which understanding the white matter may continue to have an effect on many other parts of cognitive neuroscience, even outside of the vision sciences.

Two major difficulties recur in the descriptions of findings reviewed above: the first is an ambiguity in the interpretation of findings about connectivity. In the Introduction (1), we described the Fiber Detection Problem. The FDP creates a pervasive difficulty in the interpretation of many of the findings regarding connectivity. It limits our ability to discover new tracts with tractography, and limits the interpretation of individual differences in connectivity. Changes in tractography methods or measurement parameters may reduce the missed fascicles or the false alarms, but it does not speak to increased sensitivity in detection of white matter pathways. Indeed recent reports demonstrated that increasing the resolution of the data may be only part of the story. Unless high-quality data are also associated with an optimal choice of tracking methods important and known connections can be missed (Thomas et al., 2014). The dilemma is that the best tractography analysis method is likely to depend on the properties of the data, as well as the fascicle to be estimated. For this reason it has been proposed to use personalized approaches to tracking. For the time being, no fixed set of rules will work in every case (Takemura et al., 2016). Nevertheless, finding major well-known tracts in individual humans is not a major problem, and can even be automated (Yendiki et al., 2011; Yeatman et al., 2012; Kammen et al., 2015).

The other difficulty relates to the interpretation of microstructural properties such as FA and MD. While these relate to biophysical properties of the tissue in the white matter, they contain inherent ambiguities. For example, FA may be higher in a specific location in one individual relative to another because of an increased density of myelin in that individual, but it may also be higher because of a decrease in the abundance of tracts crossing through this region. In the following last section, we outline a few directions that we believe will influence the future of the field, and perhaps help address some of these difficulties.

### Future directions

One of the common threads across the wide array of research in human vision science using dMRI that we present and a source of frustration and confusion in reading the dMRI literature is the ambiguity in interpretation: changes in parameters of models of the tissue may be caused by different biological processes. Two promising directions that we hope will address these ambiguities in the future, or at least reduce them have to do with new developments in the measurements and modeling of MRI data in human white matter:

### Biophysical models and multi b-value models

When more than one diffusion weighting b-value is collected, additional information about tissue microstructure can be derived from the data. This includes compartment models, that account for the signal as a combination of tensors (Clark and Le Bihan, 2000; Mulkern et al., 1999) or extends the Gaussian model with additional terms (so-called Diffusion Kurtosis Imaging; Jensen et al. (2005)). More recent models take into account the diffusion properties of different kinds of tissue. For example, the CHARMED model (Assaf and Basser, 2005), explicitly models intracellular water contributions to the signal and extracellular water contributions. Other models describe axon diameter distribution (Assaf et al., 2008; Alexander et al., 2010), and dispersion and density of axons and neurites (Zhang et al., 2012). However, the field is also struggling with the ambiguity of these models (Jelescu et al., 2016), and this is a very active field of research (Ferizi et al., 2016). The ultimate goal of the field in these efforts is to develop methods and models that help infer physical quantities of the tissue that are independent of the measurement device. This goal can already be achieved with other forms of MRI measurements, which we will discuss next.

### Combining diffusion-weighted and quantitative MRI

Quantitative MRI refers to an ever-growing collection of measurement methods that quantify the physical properties of neural tissue, such as their density (Mezer et al., 2013) or molecular composition (Stüber et al., 2014). These methods have already been leveraged to understand the structure and properties of the visual field maps (Sereno et al., 2013; Bridge et al., 2014). Combinations of these methods with diffusion MRI can help reduce the ambiguity and provide even more specific information about tissue properties in the white matter (Stikov et al., 2011; Mezer et al., 2013; Assaf et al., 2013; Stikov et al., 2015; Mohammadi et al., 2015). For example, a recent study used this combination to study the biological basis of amblyopia (Duan et al., 2015).

### Large open data-sets

The accumulation of large open datasets with thousands of participants will enable the creation of models of individual variability at ever-finer resolution (Pestilli and Franco, 2015). This will enable more confident inferences about the role of white matter in vision. The Human Connectome Project (Van Essen et al., 2013) is already well on its way to providing a data set encompassing more than a thousand participants, including not only high-quality dMRI data, but also measurements of fMRI of the visual system. Another large publicly available dataset that includes measurements of both dMRI and fMRI of visual areas is the Enhanced Nathan Klein Institute Rockland Sample (Nooner et al., 2012). Crucially, the aggregation of large data-sets can be scaled many times if a culture of data sharing pervades a larger portion of the research community (Gorgolewski and Poldrack, 2016) and as the technical tools that allow data-sharing become more wide-spread (Wandell et al., 2015).

**Table 1:** Publicly available data-sets for analysis of visual white matter.

| Data types | Number of available subjects | Acquisition parameters | Link | Relevant papers |
| --- | --- | --- | --- | --- |
| Human Connectome Project |  |  |  |  |
| Diffusion MRI, resting state functional MRI | 900 (as of May 2016 – eventually 1200) | b=1000, 2000, 3000, 90 directions each, 1.25 mm<sup>3</sup> isotropic resolution, 3T | http://humanconnectome.org/ | Sotiropoulos et al. (2013) |
| NKI Rockland Enhanced |  |  |  |  |
| Diffusion MRI, fMRI (including visual area localizer) | >600 (as of July 2015) | b=1500, 137 directions, 2 mm<sup>3</sup> isotropic resolution, 3T | http://fcon_1000.projects.nitrc.org/indi/enhanced/index.html | Nooner et al. (2012) |
| Data supplement to “A major human white matter pathway between dorsal and ventral visual cortex” |  |  |  |  |
| Diffusion MRI, ROIs of multiple visual field maps, tractography solutions | 5 | b=2000, 96 directions, 1.5 mm<sup>3</sup> isotropic resolution, 3T | https://purl.stanford.edu/bb060nk0241 | Takemura et al. (2015) |

### Open source software for reproducible neuroscience

One of the challenges facing researchers that are using dMRI is the diversity of methods available to analyze the data. To facilitate the adoption of methods and the comparison between different methods, another important element in progress towards reproducible science is the transparency of the methods. The combination of article text, software and data published in the notebooks that accompany this article, demonstrates how scientific concepts and findings can be communicated in writing, and augmented by the publication of computational methods that allow reproducing some or all of the computations. This contributes to the understanding of the concepts, to scientific education, the advancement of citizen science and supports computational reproducibility (Donoho, 2010; McNutt, 2014; Stodden et al., 2014; Yaffe, 2015).

To allow this reproducibility, robust and transparent open-source implementations of the methods have become crucial. There are several open-source projects focused on dMRI, and we review only a selection here (see also Table 2 for a selection of open source software for dMRI): the FMRIB Software Library (*FSL*) implements useful preprocessing tools (Andersson and Sotiropoulos, 2016), as well as individual voxel modeling (Behrens et al., 2003) and probabilistic tractography (Behrens et al., 2007). The *MRtrix* library implements novel methods for analysis of individual data, and group analysis (Tournier et al., 2012). *Vistasoft*, *mrTools*, *AFQ* and *LiFE* (Dougherty et al., 2005; Yeatman et al., 2012; Pestilli et al., 2011; Hara et al., 2014; Pestilli et al., 2014) integrate tools for analysis of visual fMRI with tools for dMRI analysis and tractography segmentation. *Camino* provides a set of tools, with a particular focus on multi b-value analysis. *Dipy* (Garyfallidis et al., 2014) capitalizes on the vibrant scientific Python community (Perez et al., 2010) and the work of the Neuroimaging in Python community (Nipy; Millman and Brett (2007)) to provide implementations of a broad array of diffusion MRI methods, ranging from newer methods for estimation of microstructural properties (Jensen and Helpern, 2010; Portegies et al., 2015; Fick et al., 2016; Özarslan et al., 2013), methods for tract clustering and tract registration Garyfallidis et al. (2012, 2015), as well methods for statistical validation of dMRI analysis (Rokem et al., 2015; Pestilli et al., 2014).

**Table 2:**
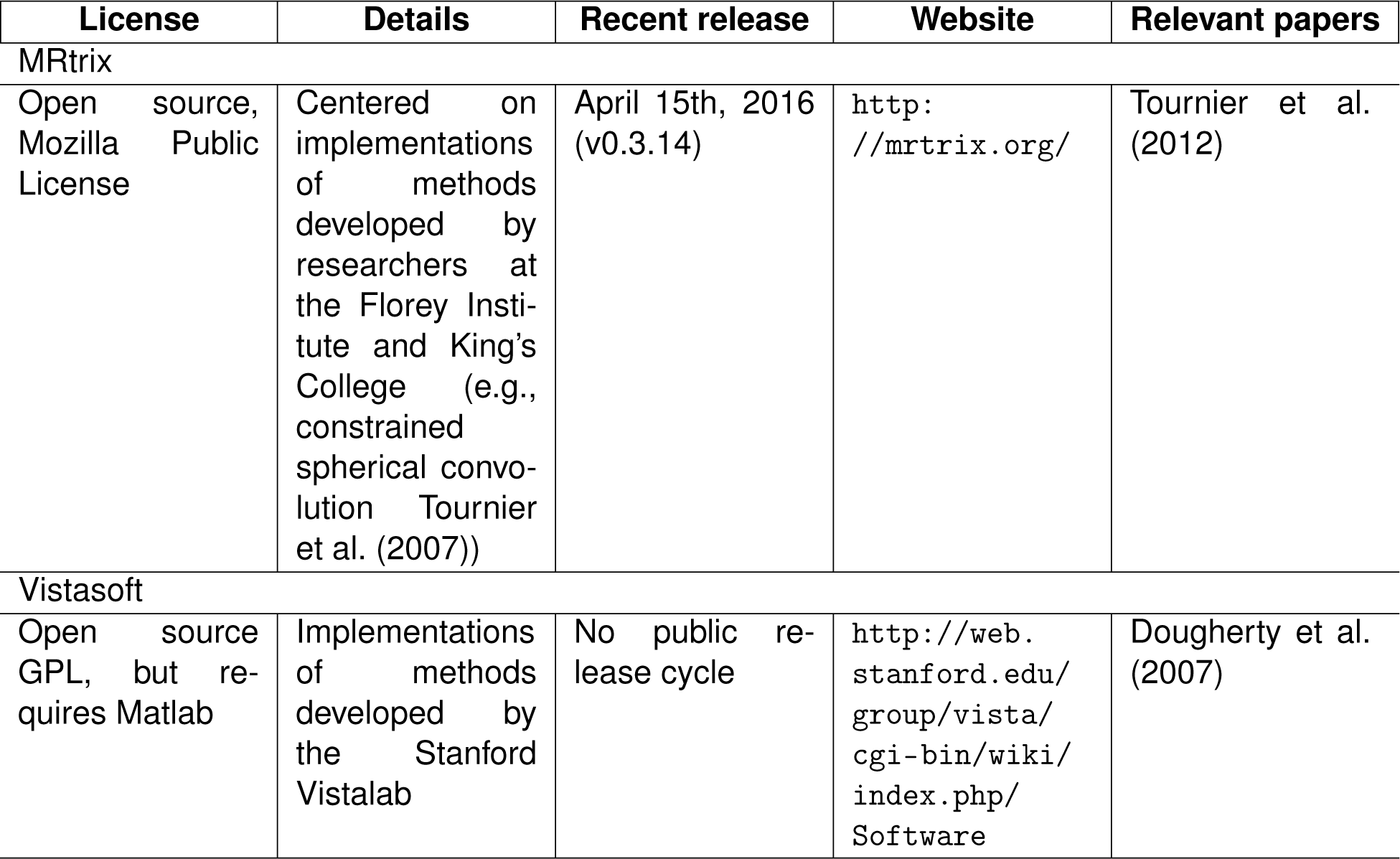
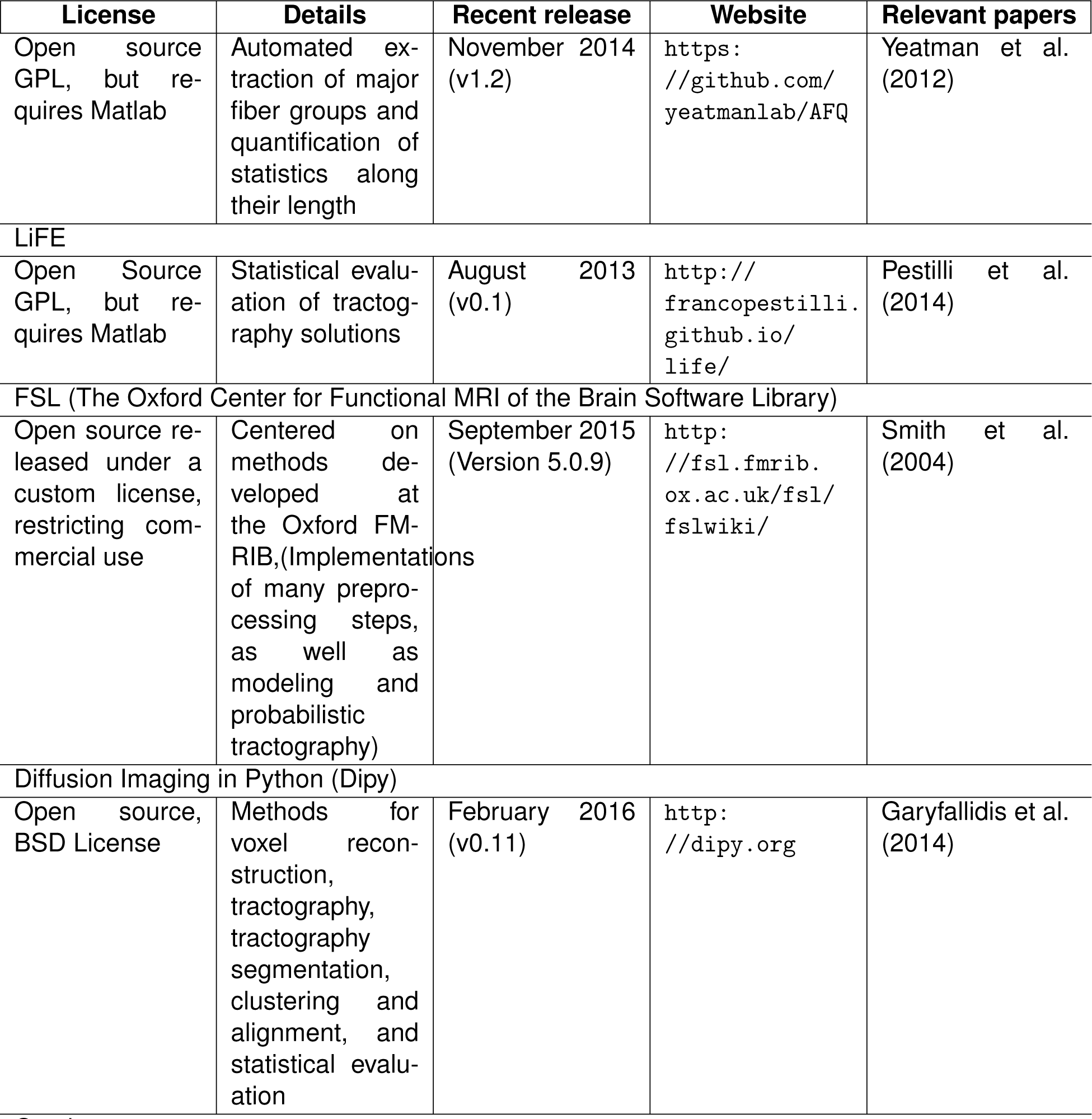
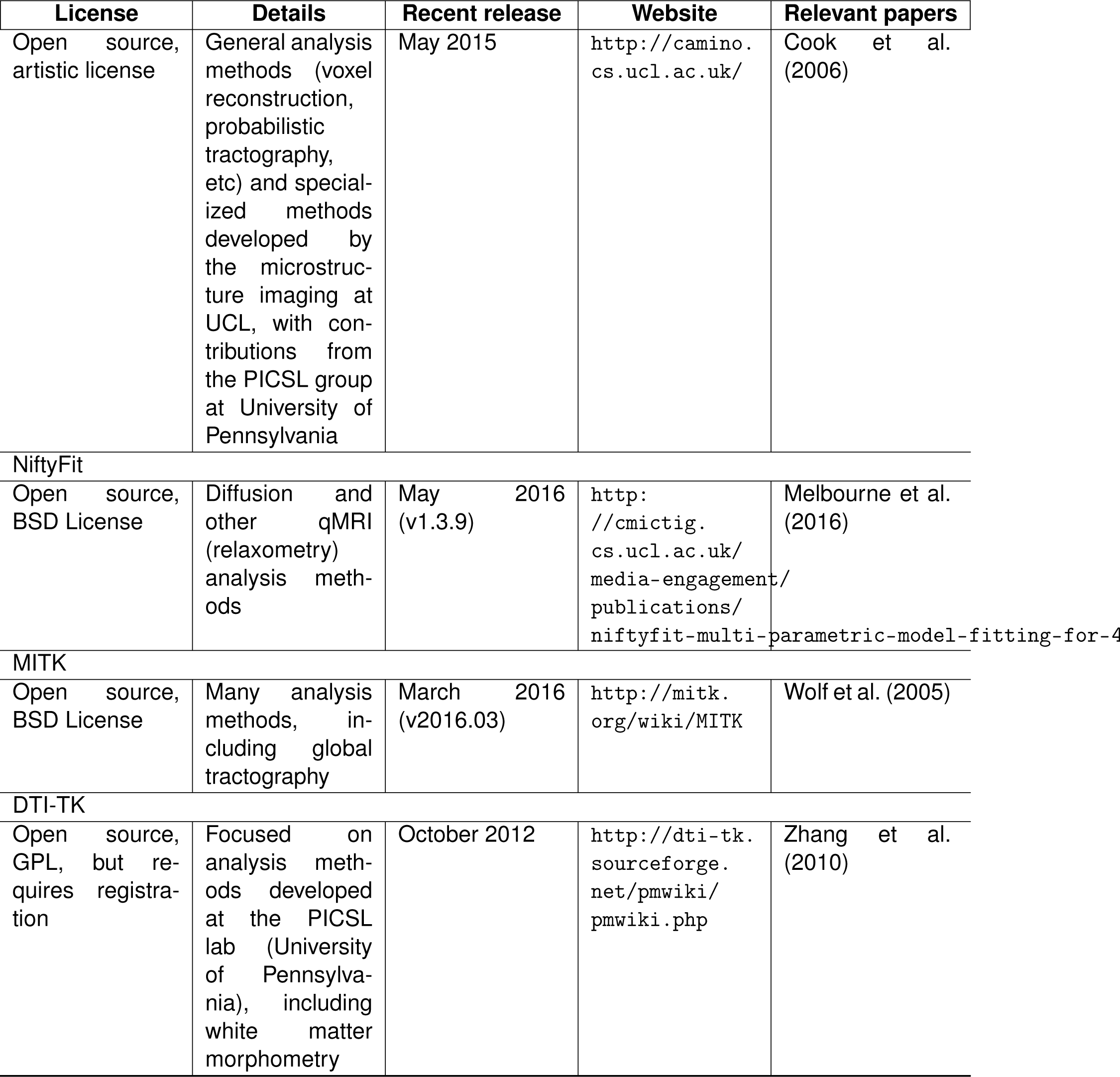
A selection of open source software for dMRI analysis.

As available data-sets grow large, another promising direction is the adoption of approaches from large-scale data analysis in neuroscience (Caiafa and Pestilli, 2015; Freeman, 2015). These methods will enable more elaborate and computationally demanding models and methods to be considered in the analysis of large, multi-participant dMRI datasets.

## Acknowledments

Ariel Rokem was funded through a grant by the Gordon & Betty Moore Foundation and the Alfred P. Sloan Foundation to the University of Washington eScience Institute Data Science Environment. Hiromasa Takemura is supported by Grant-in-Aid for JSPS Fellows. Holly Bridge is a Royal Society University Research Fellow. The work described by K. Suzanne Scherf and Marlene Behrmann was supported by Pennsylvania Department of Health SAP grant 4100047862 (M.B., K.S.S.), NICHD/NIDCD P01/U19 (M.B., PI-Nancy Minshew), NSF grant (BCS0923763) to M.B., National Institutes of Mental Health (MH54246) to M. B., NSF Science of Learning Center grant, “Temporal Dynamics of Learning Center” (PI: Gary Cottrell, Co-I: M.B.), a post-doctoral fellowship from the National Alliance for Autism Research to K.S.S. and Beatriz Luna, and by awards from the National Alliance of Autism Research to Cibu Thomas and Katherine Humphreys and from the Cure Autism Now foundation to Katherine Humphreys. Work described by Ione Fine, Holly Bridge and Andrew Bock was funded by the National Institutes of Health (EY-014645); Holly Bridge is a Royal Society University Research Fellow. Franco Pestilli was funded by the Indiana University College of Arts and Sciences, the Indiana Clinical and Translational Institute (NIH UL1-TR001108). We thank the Jupyter team at UC Berkeley and beyond for creating the notebook format that is used in the example code provided as a computational appendix, and the Freeman Lab at HHMI Janelia Research Campus for creating the “Binder” platform to run these notebooks. We thank Kendrick Kay, Brent McPherson, Daniel Bullock, Sophia Vinci-Booher, Sandra Hanekamp, Shiloh Cooper, Brian Allen, Ian Chavez for comments on early versions of the manuscript.

